# kmerRRR: A k-mer based tool for functional genomics in Repeat Rich Regions

**DOI:** 10.64898/2026.06.21.732238

**Authors:** Jabale Rahmat, Tuan Pham, Amanda M. Larracuente

**Affiliations:** Department of Biology, University of Rochester, Rochester, NY, USA

## Abstract

Highly repetitive sequences pose problems for genome assembly and analysis. While advances in long-read sequencing technologies have helped reveal the organization of repetitive genomic sequences at unprecedented resolution, their functional characterization remains difficult because molecular assays that probe protein-DNA interactions and characterize expression often rely on short read sequencing. The repetitive nature of these regions poses major challenges for methods relying on sequence mapping, which is exacerbated for short reads. Repetitive genome regions often have low mappability, leading to substantial information loss during downstream filtering. To address this challenge, we developed a bioinformatic tool—kmerRRR—that leverages k-mer frequency analyses to enhance the mappability of repetitive regions. KmerRRR compares k-mer frequencies within user-defined loci to their frequencies across the genome to identify repetitive sequences that are overrepresented locally relative to the global background. This approach quantifies locus uniqueness, allowing users to distinguish sequences that are globally repetitive from those that are repetitive, but restricted to specific genomic loci. We demonstrated the utility of this method by reanalyzing chromatin profiling data from human, *Drosophila*, and *Arabidopsis* centromeres and small RNA sequencing data. Our results show that incorporating local k-mer ratio information enhances read retention and signal interpretation within repetitive regions, thereby recovering biologically meaningful information that is typically lost in conventional analyses. The tool is freely available under MIT license in github: (https://github.com/LarracuenteLab/kmerRRR).

## Introduction

Heterochromatic regions of the genome are generally gene-poor and repeat-rich: the bulk of these regions consists of tandem repeats and transposable elements (TEs). These highly repetitive genome regions are important to understand, as they can contain essential chromosomal domains like centromeres (Brar & Amon, 2008), telomeres (Di Pietro et al., 2024), and ribosomal DNAs (Di Pietro et al., 2024). Tandemly-repeated sequences pose problems for genome assembly and analysis. However, recent advances in long-read sequencing technology (Wenger et al., 2019) and assembly algorithms (Cheng et al., 2021; Cheng et al., 2026; Rautiainen et al., 2023) have dramatically improved our ability to resolve the organization of repeat-rich genome regions ((Naish et al., 2021; Nurk et al., 2022), reviewed in (Li & Durbin, 2024)). This includes large arrays of satellite DNAs (satDNAs), which are among the most repetitive sequences in eukaryotic genomes and are often associated with centromeres, telomeres, and sex chromosomes. However, understanding the functional genome features and properties associated with repetitive DNA remains challenging.

Repetitive DNA sequences can make up a large fraction of eukaryotic genomes. They play roles in genome organization (Jagannathan et al., 2018; Jagannathan et al., 2019; Jagannathan & Yamashita, 2021) and genomic stability (e.g. (Flynn & Yamashita, 2024)), chromosome segregation (e.g. (Altemose, Logsdon, et al., 2022)), and heterochromatin formation (e.g. (Wei et al., 2021)). Genomic instability associated with highly repetitive DNA is implicated in multiple human diseases, including cancer (reviewed in (Ugarković et al., 2022)).

Despite increasing evidence that highly repetitive sequences influence important cellular processes, the mechanisms underlying many of these functions are poorly understood. Addressing these questions requires functional genomic approaches capable of interrogating highly repetitive genomic regions.

To gain deeper insights into functional features of repetitive genome regions requires complementary functional genomic approaches such as those that detect transcription factor binding, chromatin state, transcription, and genetic variation. Many of these approaches remain dependent on short-read sequencing (Park, 2009; Skene & Henikoff, 2017), presenting a challenge for the analysis of highly repetitive genomic regions (Park et al., 2025). Although k-mer based analyses have helped alleviate some assembly challenges (reviewed in (Moeckel et al., 2024)) and have proven useful for characterizing satDNA arrays (Wei et al., 2014) and assessing polymorphism among individuals in human populations (Altemose et al., 2014), accurately mapping short reads to distinct genomic loci remains challenging.

For reference-based analysis approaches, researchers make functional genomic inferences based on how reads map back to the genome. A typical step in these genomic analyses involves filtering reads by their mapping quality (MAPQ): Reads that map uniquely are assigned high confidence MAPQ scores by alignment tools (Almeida da Paz et al., 2024; Langmead & Salzberg, 2012; Li, 2018; Li & Durbin, 2009). Reads originating from repetitive regions, containing sequencing errors, or representing chimeric fragments often map to multiple genomic locations and therefore receive low MAPQ scores (Almeida da Paz et al., 2024). Such low-confidence reads are frequently filtered out during downstream analyses. For highly repetitive sequences that are widely distributed across the genome, this strategy is warranted and acknowledges a limitation of reference-based genome analysis.

This approach to quality control treats the majority of reads mapping to repetitive regions as largely uninformative. However, some tandem repeats may have restricted genomic distributions where sequence reads will map multiply but are still informative about the genomic locus despite their low mappability. We refer to these sequences as locus-unique: they are repetitive, occurring multiple times within a restricted genomic region, but rarely elsewhere in the genome. Existing methods to summarize mappability consider uniqueness across the whole genome, leading to low mappability in these regions (Almeida da Paz et al., 2024; Karimzadeh et al., 2018; Pockrandt et al., 2020). Standard MAPQ filtering approaches do not distinguish these locally mappable reads from those mapping to repeats broadly distributed across the genome, resulting in the loss of potentially informative reads (Almeida da Paz et al., 2024). As a result, biologically relevant signatures can be overlooked, limiting the ability to infer functional genomic activity in repetitive genome regions.

To address this limitation, we developed kmerRRR, a k-mer-based framework that quantifies local mappability directly from genome assemblies. By comparing local and genome-wide k-mer frequencies, kmerRRR identifies sequences that are repetitive, but locus-unique within user-defined regions. KmerRRR also provides the option to incorporate this local mappability information into downstream analyses and retain informative reads that would otherwise be discarded by standard MAPQ-based filtering. Although biologically meaningful application of kmer requires a well-annotated genome assembly or clearly defined genomic regions of interest, the framework enables functional genomic analyses in regions that are often excluded from conventional workflows.

## Methods

### kmerRRR algorithm and LUC inference

Figure 1 summarizes the core workflow of kmerRRR. The central module, *kmers_stat* infers local mappability for user-defined loci provided in BED format. For a specified k-mer length *k*, the genome-wide (global) k-mer counts are computed first and then the locus-specific (local) k-mer counts for each locus in the BED file is computed. For every k-mer within a locus, a local-to-global ratio is calculated, generating an intermediate table of per k-mer enrichment statistics. To infer local mappability at nucleotide resolution, each locus is scanned with a step size of 1. At each base position *j*, all overlapping k-mers are collected, and their counts and local-to-global ratios are aggregated. The total k-mer support per base is defined as the sum of counts of all overlapping k-mers. In parallel, summary statistics of per-base local-to-global ratios (total, mean, median, mode, maximum, and minimum) are computed. Locus-Unique Characters (LUCs) are then inferred using two criteria: *i*) a repeat support criterion based on total k-mer counts; and *ii*) a ratio enrichment criterion based on the mean local-to-global ratio exceeding a user-defined threshold. The combination of these criteria enables classification of each nucleotide as globally unique, repetitive but locus-unique, globally repetitive, or inconclusive. The repetitive flag distinguishes repetitive from non-repetitive bases, while l_unique Identifies both locus-unique and globally unique positions. These are subsequently parsed to differentiate locus-unique repetitive bases from globally unique regions.

**Fig1.**
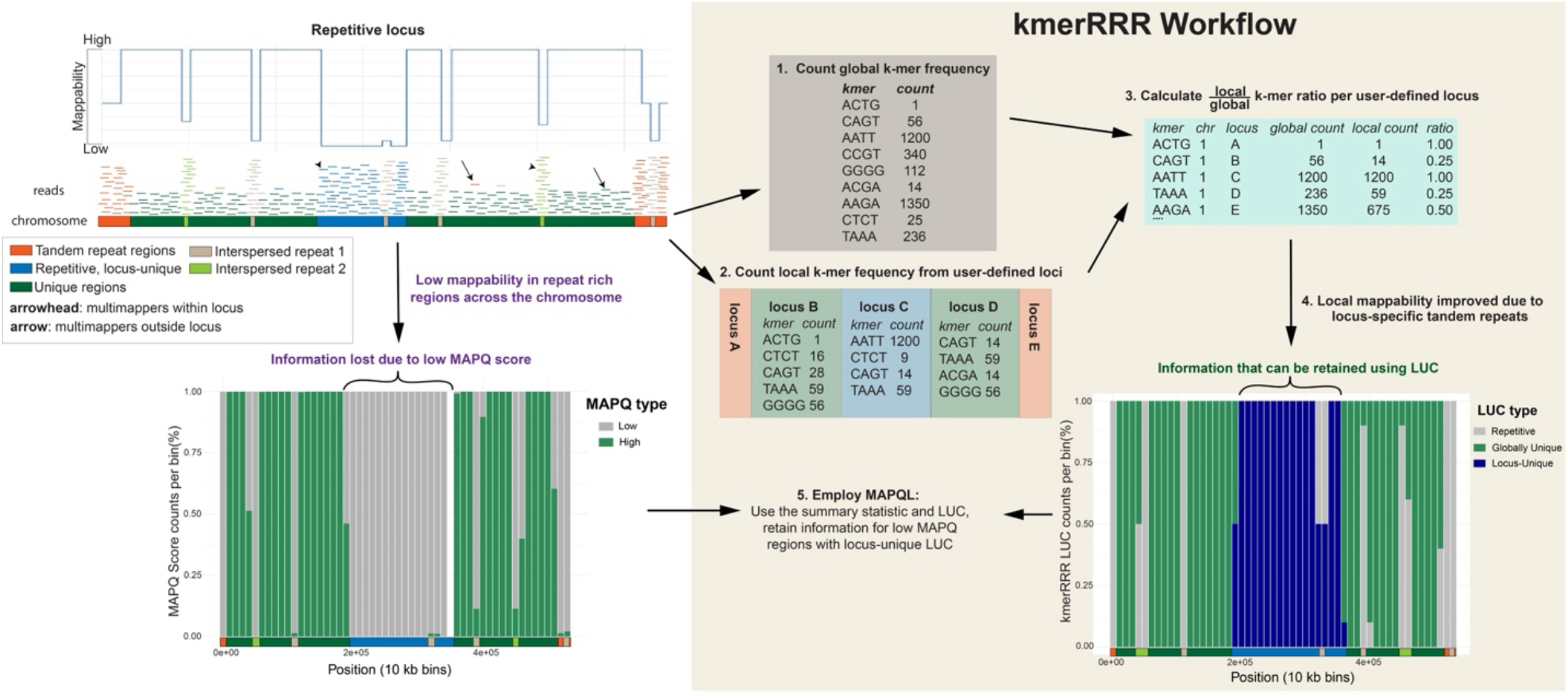
Schematic overview of the kmerRRR workflow. Repetitive regions create mappability challenges across a simulated chromosome. Short reads originating from repeat-rich sequences frequently generate multi-mapping alignments, resulting in ambiguous placement and low mapping quality (MAPQ) scores. kmerRRR addresses this limitation by leveraging genome wide k-mer frequency information to compute local-to-global k-mer ratios within user-defined loci and by inferring Locus-Unique Character (LUC) at nucleotide resolution using k-mer abundance and local-global ratio statistics. This framework identifies regions that are globally repetitive but locus-unique, thereby improving locus-specific mappability. The resulting mappability landscape, together with inferred sequence context, enables retention of informative reads in BAM files that would otherwise be discarded, enhancing downstream functional analyses in repeat-rich regions.

Pseudocode

Let:

G(k) = global k-mer count table

L_i_(k) = local k-mer count table for locus *i*

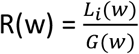 = local-to-global ratio for k-mer *w*

µ_j_ = mean{R(w) : w ∈ B_j_}

B_j_ = set of k-mers overlapping base position *j*

T_j_ = ∑w∈*B*_j_ *G*(*w*) (or local counts, depending on implementation)

User-defined parameters:

τ_*c*_ = minimum cumulative k-mer support threshold at a position (Cumulative support is defined as the total count of all k-mers overlapping a given nucleotide position. Positions near the ends of a sequence are overlapped by fewer k-mers than positions in the sequence interior, where the maximum number of overlapping k-mers equals the k-mer length, *k*.)

*τ*_*r*_ = minimum mean ratio threshold

~~~
For each locus i in BED:
  For each base position j in locus i:
    B_j ← all k-mers overlapping j
    T_j ← sum of k-mer counts in B_j
    µ_j_ ← mean local-to-global ratio in B_j
    if T_j > *τ*_*c*_:
        repeat_quality ← 1
    else:
        repeat_quality ← 0
    if µ_j_ ≥ *τ*_*r*_:
        ratio_quality ← 1
    else:
        ratio_quality ← 0
    if repeat_quality == 1 and ratio+quality ==1:
        repetitive ← 1
        l_unique ← 1
    elif repeat_quality == 0 and ratio+quality ==1:
        repetitive ← 0
        l_unique ← 1
    elif repeat_quality == 1 and ratio+quality ==0:
        repetitive ← 1
        l_unique ← 0
    else:
        repetitive ← 0
        l_unique ← 0
~~~

#### Post processing step

If l_unique = 1 and repetitive = 1 → locus-unique

If l_unique = 1 and repetitive = 0 → globally unique

If l_unique = 0 and repetitive = 1 → globally repetitive

Else → inconclusive

#### Mathematical clarification of thresholds

To formalize the criteria more rigorously:

For base position *j* within locus *i*

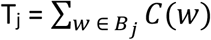

Where C(w) is the chosen k-mer count metric (global or local, depending on implementation).

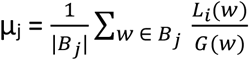

A base is classified as:

repetitive, if T_j_ > τ_*c*_

locus-unique, if μ _j_ ≥ τ_*r*_

where, *τ*_*c*_ controls sensitivity to repetitive support and *τ*_*r*_ controls stringency of local enrichment. This formulation makes explicit that LUC inference depends both on abundance (count support) and relative enrichment (local/global ratio), allowing discrimination between globally unique, locus-unique, and globally repetitive sequence contexts.

### Application of LUCs: Retaining multi-mapping reads from BAM files

To extend the utility of LUCs to downstream analyses, kmerRRR includes a *bam_file_manipulate* script that adjusts mapping quality scores based on local mappability information. Specifically, the script assigns a modified mapping quality score, termed MAPQL (Mapping Quality based on local mappability), to reads overlapping locus-unique regions. Importantly, MAPQL is not recalculated from the underlying alignment algorithm. Instead, *bam_file_manipulate* leverages LUC annotations inferred by *kmers_stat* in combination with the existing alignments present in the BAM file. For loci defined in the user-provided BED file, the script scans reads mapped to the targeted chromosome and selectively modifies reads with the low MAPQ scores that would otherwise be discarded during standard filtering.

For each read, the mean per base local-to-global k-mer ratio across the aligned region in computed as:

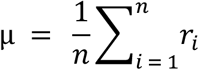

where r_i_ represents the mean k-mer ratio at base position *i* and *n* is the number of positions spanned by the read. If the computed mean ratio exceeds a user-defined threshold, and the original MAPQ is below a specified cutoff, the script updates the mapping quality to MAPQL, allowing the read to be retained. Reads that are unmapped or hard-clipped are automatically excluded from manipulation. For computational efficiency, we recommend that users provide pre-filtered from all criteria except mapping quality. This reduced processing time while ensuring that only reads subject to MAPQ based exclusion are evaluated for potential retention.

Because MAPQ scores are traditionally interpreted as probabilities of correct alignment, modifying them could raise concerns about artificially inflating mapping confidence. However, MAPQL does not alter the underlying alignment coordinates, CIGAR string, or alignment score. Instead, it reevaluates mapping confidence in the context of locus-specific uniqueness derived from genome-wide k-mer distributions.

Conventional aligners compute MAPQ based on global repetitiveness, penalizing reads originating from repetitive arrays even when those arrays are only repetitive within a defined locus (e.g., a specific centromere). kmerRRR addresses this limitation by incorporating independent sequence-context information derived from local-to-global k-mer ratios. Reads are only retained if their mean per-base local enrichment exceeds a user-defined threshold, ensuring that rescued reads originate from regions with demonstrable locus specificity.

Importantly, reads are modified only if they overlap locus-unique bases (LUCs), their mean local-to-global ratio exceeds a defined threshold, and their original MAPQ falls below the filtering cutoff. In this process, no new alignments are introduced, no ambiguous multi-locus reads are reassigned, or no coordinate changes occur. Thus, MAPQL should be interpreted not as a replacement for alignment-based MAPQ, but as an orthogonal confidence metric reflecting locus-specific mappability within repetitive regions. The reads that are adjusted with MAPQL are explicitly annotated in the BAM file with the tag “KR:LUC” in the read group field.

To avoid incorporating duplicate reads during BAM file manipulation, kmerRRR randomly selects among reads that satisfy the filtering criteria using Python 3’s random module. Seed values are generated from operating system entropy and recorded in the log file, allowing analyses to be reproduced when the same seed is supplied. Consequently, manipulated BAM files may vary slightly between runs, although the resulting estimates and biological conclusions are expected to remain unchanged.

### Software implementation and dependencies

For k-mer counting, kmerRRR incorporates Jellyfish (Marcais & Kingsford, 2011), a fast memory efficient k-mer counting tool. The program invokes the Jellyfish through Python 3 using the *subprocess* module to execute counting commands. To query and access the resulting k-mer databases, it uses Pyjellyfish, a Python wrapper that enables efficient retrieval of k-mer counts within the analysis pipeline.

Genome sequence parsing is performed using the *SeqIO* module from Biopython, which facilitates efficient reading and handling genome sequence files. For BAM file processing, read filtering, and MAPQ manipulation, it uses pysam, a Python interface to SAMtools (Danecek et al., 2021) to enable programmatic access to alignment files.

The kmerRRR framework is implemented in Python 3 and leverages built-in utilities for genomic interval parsing and file operations. Numerical computations and summary statistics (e.g., mean, mode, minimum, and maximum k-mer ratios) are computed using the NumPy package to ensure computational efficiency and numerical robustness.

In addition to the jellyfish-based implementation, an alternative k-mer counting module using Python dictionaries is developed. While this option provides portability and simplicity for small-scale analyses, it is recommended that the jellyfish-based workflow for genome-scale datasets should be used due to its superior performance in runtime and memory efficiency.

KmerRRR includes plotting functionality implemented in R (version ≥ 4.3) (R Core Team, 2005). These scripts are executed from Python 3 using the subprocess module (see documentation).

### Genome analysis

For genome analyses, human T2T-CHM13v2.0 genome from the github page: https://github.com/marbl/CHM13, and *Arabidopsis thaliana* near T2T genome from the github page: https://github.com/schatzlab/Col-CEN/tree/main/v1.2 were used. For the human genome, the mitochondrial sequence was excluded from all our analyses. The *D. melanogaster* genome was analyzed using the assembly generated by Chang & Larracuente (2019). Centromere loci for *H. sapiens, A. thaliana*, and *D. melanogaster*, were annotated according to Altemose *et al*. (2022), and Naish *et al*. (2022), and Chang *et al* (2019), respectively.

### ChIPseq and CUT&TAG analysis

The datasets used in this study are in Table 1. Each dataset was downloaded using sratoolkit (https://github.com/ncbi/sra-tools) version 3.0.0 with *prefetch* followed by *fasterq-dump* with *--split-files --threads 8* parameters. For each FASTQ files, the adapters were removed using trimgalore version 0.6.2 (Krueger, 2026) using *trim_galore --paired --phred33 --fastqc --length 35* options. Bowtie2 version 2.3.5.1 (Langmead & Salzberg, 2012) with *bowtie2 -x index -1 reads1*.*fq -2 reads2*.*fq --very-sensitive -q -S output*.*sam* options was used for mapping the paired-end reads to respective reference genomes. For bwa version 0.7.17 (Li & Durbin, 2009), *bwa mem -t 32 -a ref*.*fasta reads1*.*fq reads2*.*fq > output*.*sam* was used for mapping the CUT&Tag reads to the respective reference genomes. For converting SAM files to BAM files and filtering, samtools version 1.21 (Danecek et al., 2021) was used with *samtools view -f 3 -F 4 -F 8 -F 256 -F 2048 -q 30* options. Peaks were called using MACS version 2.1.1.20160309 (Feng et al., 2012) with *macs2 callpeak -t treatment*.*bam -c control*.*bam -f BAMPE -g (dm, hs, or 1e8 for D. melanogaster, Homo sapiens, and A. thaliana*, respectively*) -q 0*.*01 -n output -B --call-summits --outdir output*.*directory* options. For evaluating overlapping peaks, idr version 2.0.3 (Li et al., 2011) was used with *idr --samples peak1*.*narrowPeak peak2*.*narrowPeak --input-file-type narrowPeak --output-file-type narrowPeak --rank p*.*value -l file*.*log -o peaks*.*idr* after sorting the files using *sort -k8,8nr* for p-values.

**Table 1.**
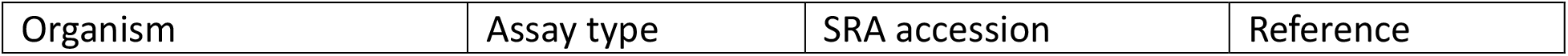
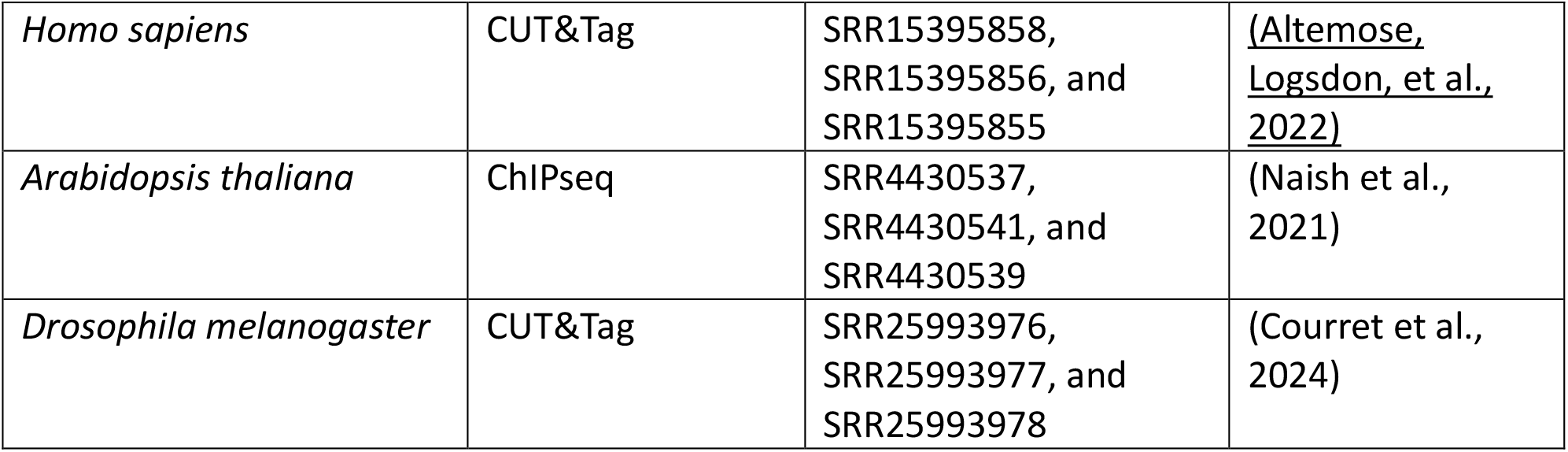
SRA numbers for CUT&TAG and ChIPseq analyses used for this study.

## Results

### Distribution of locus-vs global mappability

By leveraging local vs global k-mer frequency ratios, kmerRRR generates high resolution local mappability profiles across chromosomes. We applied this approach to the T2T *Arabidopsis thaliana* genome (Naish et al., 2021) and to chromosomes 9 and 22 of the human T2T genome (Altemose, Logsdon, et al., 2022; Nurk et al., 2022). Using 21-, 31-, 61-, and 151-mers, we characterized the distribution of Locus-Unique Characters (LUCs) across each chromosome at 100-kbp windows while keeping the centromere as a separate locus of its own and compared these profiles to traditional mappability estimates obtained by remapping all possible sequences of corresponding lengths of *k* back to the genome (Fig 2 and Supplemental Fig S1 & S2).

**Fig2.**
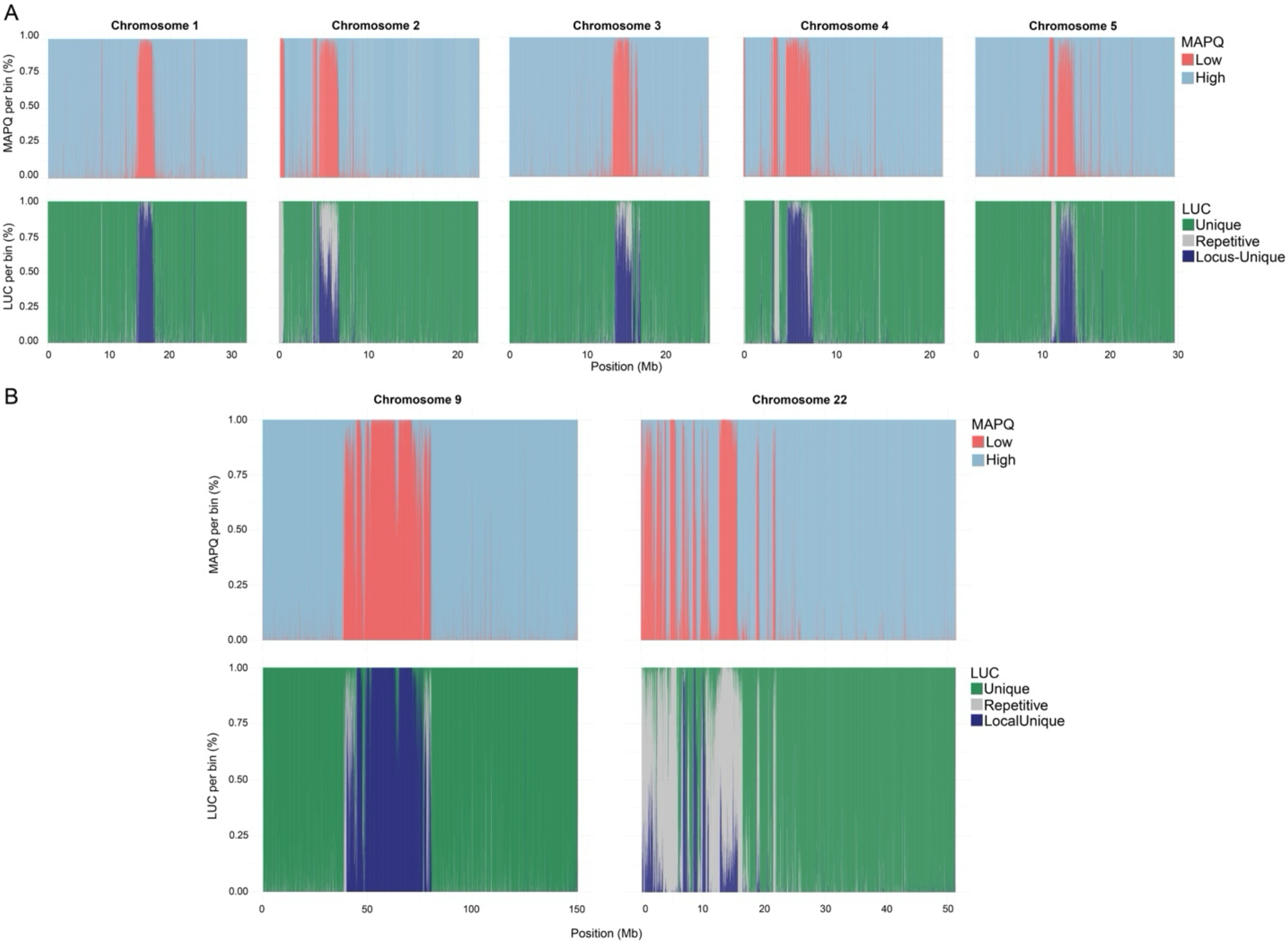
Genome wide mappability distribution in the T2T assemblies. (A) *A*.*thaliana* near T2T genome and (B) *H. sapiens* T2T chromosome 9 and 22. Each chromosome was fragmented into 150bp sliding window with a step size of 1bp, and all resulting sequences were mapped back to their respective genomes. The top panels showing the distribution of MAPQ scores across each chromosome for all 150-bp placements. Repetitive regions predominantly corresponding to centromeric arrays, exhibit low MAPQ scores, reflecting multi-mapping ambiguity. kmerRRR was applied using 151-mers, with loci defined as 100-kbp windows across each chromosome, while centromeric regions were treated as single independent loci. The lower panels illustrate the inferred locus-unique character (LUC) landscape across each chromosome inferred using a local-to-global k-mer ratio of 1. Regions that are globally repetitive but local unique show improved locus-level mappability under the kmerRRR framework.

We generated local mappability profiles across each chromosome of *A. thaliana* by partitioning the genome into 100-kbp windows and defining the centromere as an independent locus (Fig 2 and Supplemental Fig S1 & S2)(Naish et al., 2021). As expected, increasing k-mer length improves the accuracy of local mappability inference while reducing overall genomic coverage, enriching locus-unique signal at specific loci (Fig2 and Supplemental Fig S1). Accordingly, the number of inferred locus-unique bases increased from 2,968,664 genome-wide with 31-mers, including 545,504 centromeric bases, to 10,942,465 genome-wide with 151-mers, including 8,401,411 centromeric bases. kmerRRR could infer the local mappability within centromeric repeat arrays in addition to interspersed repeat contents across the chromosomes (Fig 2 and Supplemental Fig S1-S3). At smaller *k* values, k-mers are more likely to match consensus repeat sequences shared across chromosomes, making them locus non-specific. This is reflected in reduced “locus-unique” LUC inference at lower *k* values (Supplemental Fig S1). For benchmarking kmerRRR, we have used *k* up to 151, although users may choose larger values, depending on repeat monomer length. However, increasing *k* comes with a trade-off for greater storage capacity and larger RAM space. For example, when kmerRRR is applied to all centromeric regions of the human genome, increasing the k-mer size from 31 to 151 doubles storage requirements (~50GB to ~100GB) and increases memory usage from approximately 150GB to 400GB, highlighting the computational costs associated with larger k-mer sizes. In addition, the usage of local-to-global mean k-mer ratio threshold is important in inferring LUCs. We have used a mean threshold of 1 and 0.95 per nucleotide (Supplemental Fig S1 & S2).

Increasing this mean ratio threshold makes inference more stringent and improves specificity, whereas lower thresholds identify more locus-unique LUCs at the cost of reduced accuracy (Supplemental Fig S2), although depending on the context, the threshold can be any value between 0 and 1.

Because k-mer counting typically includes both DNA strands, we used canonical k-mers, which is defined as the lexicographically smaller sequence between a k-mer and its reverse complement, to avoid strand-specific redundancy. Switching from canonical to non-canonical counting altered LUC inference at smaller *k* values (~12-fold increase in centromeric locus-unique positions, from 545,504 using canonical 31-mers to 6,434,179 using non-canonical 31-mers), whereas larger *k* values showed no substantial differences (~1.1-fold increase, from 8,401,411 to 9,276,958 centromeric locus-unique positions identified using canonical and non-canonical 151-mers, respectively; Supplemental Fig S3).

Interestingly, within centromeric loci, we observed interspersed chromosome-specific unique segments, potentially arising from chromosome-specific sequence divergence or TE insertions that create unique junctions between centromeric satDNA and TEs (Fig2 and Supplemental Fig S7).

To evaluate the distribution of local mappability across the genome, we simulated 150-bp paired-end Illumina HiSeq 2500 reads using ART (Huang et al., 2012) and mapped the simulated reads back to their respective genomes. As expected, repetitive regions exhibited low MAPQ scores under standard alignment, reflecting their global repetitiveness (Fig 3 and Supplemental Fig S4 & S5). We then incorporated LUC derived from local mappability information from kmerRRR and adjusted MAPQ scores based on local k-mer enrichment (MAPQL, Fig3 and Supplemental Fig S4 & S5). Repetitive regions that were locus-unique showed a substantial increase in MAPQ, significantly improving their effective mappability without introducing detectable false positives or false negatives.

**Fig3.**
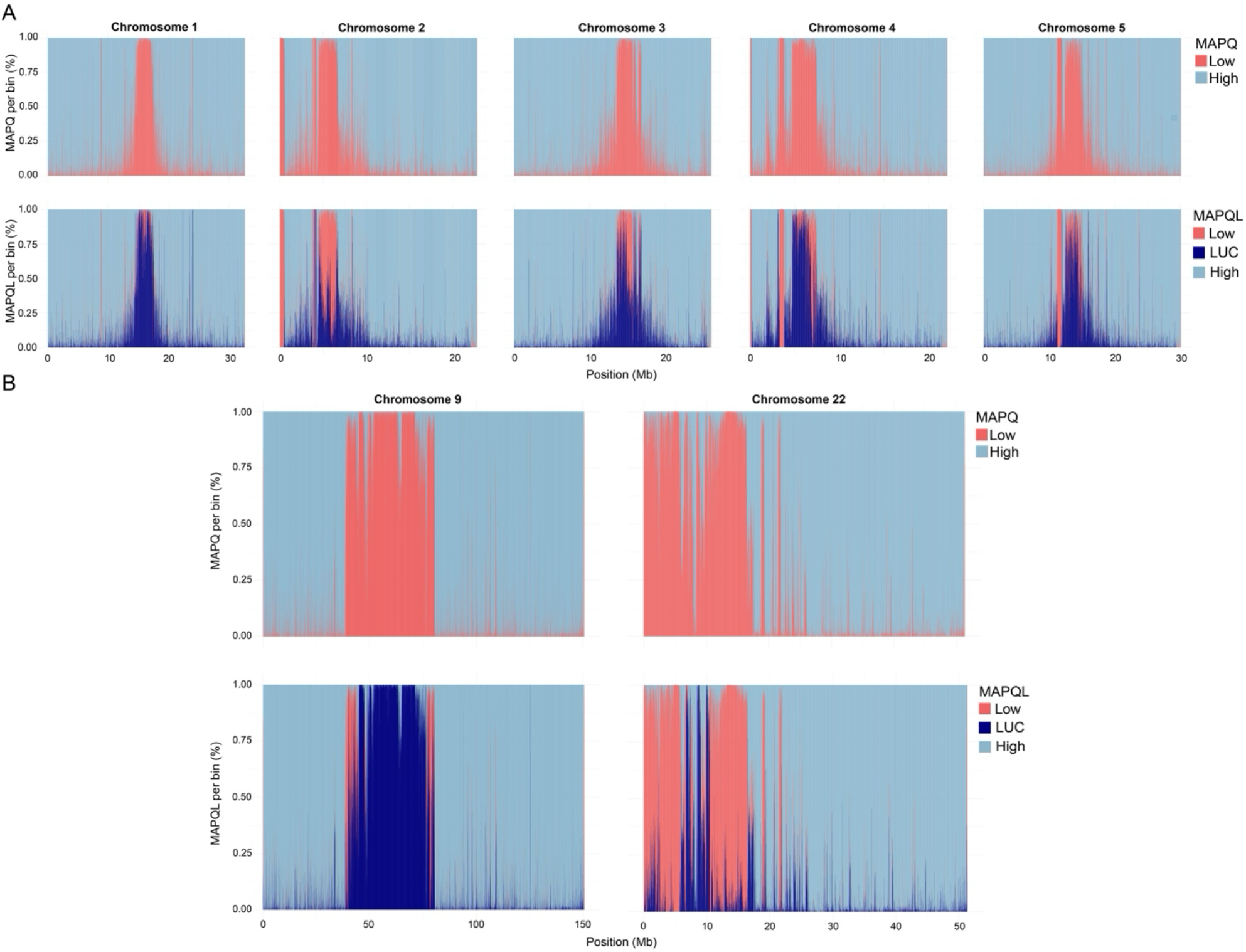
kmerRRR improves local mappability in simulated reads. MAPQ distribution across *A. thaliana* near T2T genome (A) and Human T2T genome (B) using paired-end simulated illumina sequence reads, shown before and after incorporating kmerRRR based local mappability adjustment. Highly repetitive regions exhibit low MAPQ scores under standard alignment (top panels). Integration of local mappabiliy information derived from local to global k-mer ratios increases mapping quality in regions with high k-mer ratios (MAPQL), indicating locus-unique segments that would otherwise be discarded (bottom panels).

We have also incorporated a script in kmerRRR that can create bedGraph format files, enabling direct visualization in genome browsers such as Integrative genomic Viewer (IGV) (Robinson et al., 2011) for integration with genome annotations and other genome tracks (Supplemental Fig S6). To demonstrate the utility of bedGraph files, we used the *A. thaliana* genome as an example (Fig S6). BedGraph tracks facilitate exploration of local mappability patterns in IGV, enabling both locus-specific inspection and genome-wide visualization from a broader perspective (Fig S6).

Together, these results demonstrated that kmerRRR leverages genome-wide k-mer frequency distributions to distinguish locus-unique repetitive regions from repeats that are broadly distributed across the genome. This framework enhances the interpretation of local mappability within large repetitive arrays by identifying sequences that are locus-unique even when they are globally repetitive.

#### Leveraging local mappability to enhance read retention and peak calling

Repetitive elements are often associated with centromeric regions. To investigate centromere sequence composition and infer centromeric protein localization, short read datasets from ChIPseq, CUT&Tag, and CUT&RUN experiments are usually mapped back to the reference genome (Kaya-Okur et al., 2019; Park et al., 2025; Park, 2009; Skene & Henikoff, 2017). However, because these datasets consist of short reads, downstream analyses in repetitive regions are challenging and often require stringent MAPQ filtering, which can lead to substantial loss of biologically informative reads (Supplemental Fig S9).

To evaluate the utility of local mappability inferred by kmerRRR, we used per base local to global k-mer ratios to selectively retain reads from BAM files that would otherwise be discarded due to low MAPQ scores. We applied this approach to centromeric protein profiling datasets from *A. thaliana* (ChIP-seq), *H. sapiens* (CUT&Tag), and *D. melanogaster* (CUT&Tag) (Altemose, Logsdon, et al., 2022; Chang et al., 2019; Naish et al., 2021).

Using local mappability information (Supplemental Fig S8), we retained additional reads in repetitive centromeric regions without introducing detectable erroneous mappings, thereby improving inference of CENP-A enrichment profiles (Fig4 and Supplemental Fig S9). Across all treatment and control datasets, the total number of reads passing MAPQ based filtering increases after incorporating kmerRRR based local mappability. Furthermore, peak profiles became more contiguous across centromeric loci for all the datasets (Fig 4).

**Fig4.**
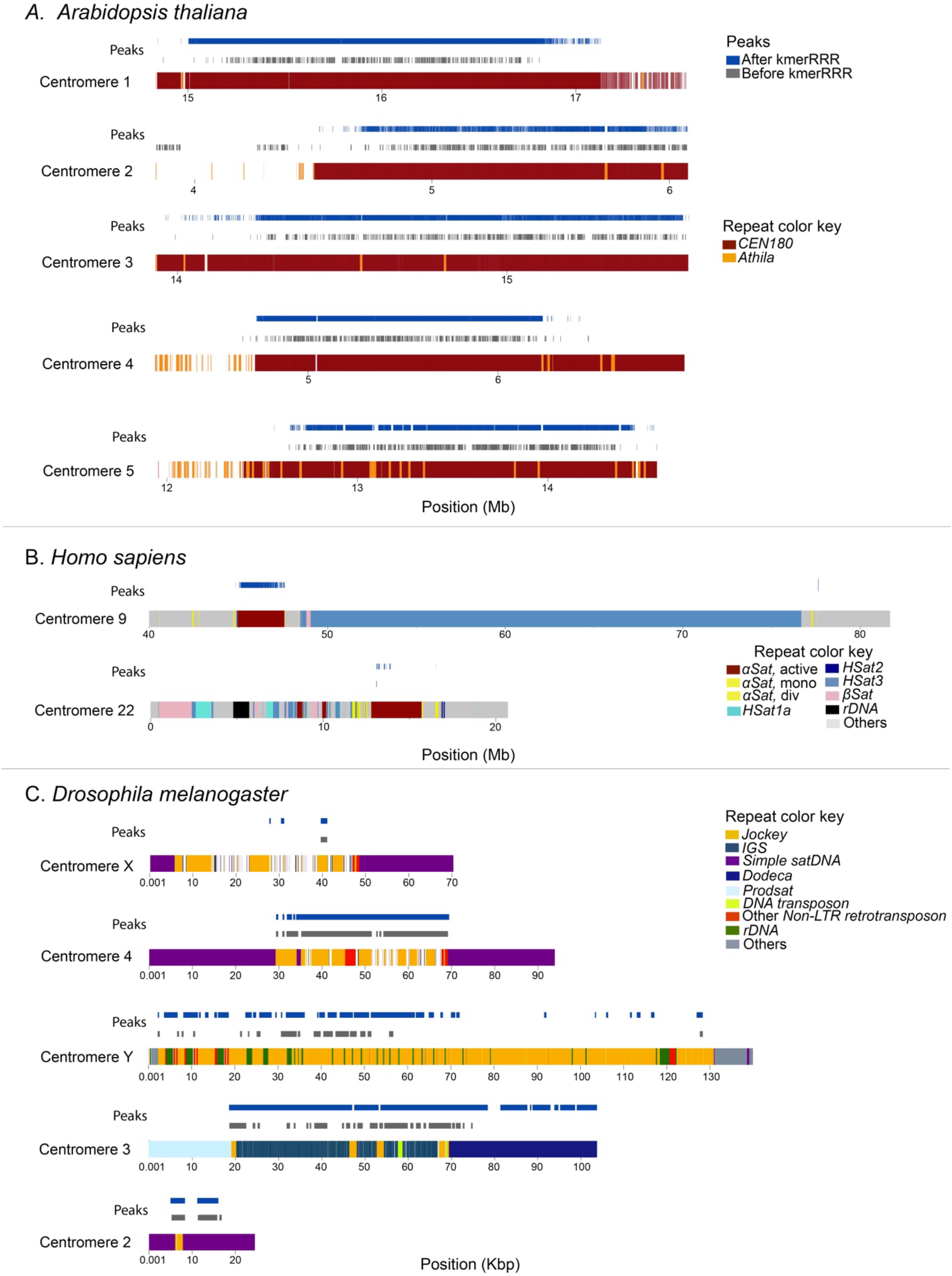
kmerRRR enhances peak calling in centromeric regions from ChIP-seq and CUT&Tag datasets. Peak profiles across centromeric regions of *A. thaliana* (A), *H. sapiens* (B), and *D. melanogaster* (C) before and after incorporating kmerRRR derived local mappability information. Prior to integrating local to global k-mer ratios, peaks were sparse and frequently absent in regions with high repeat content. Retaining reads from locus-unique segments based on k-mer ratio substantially improves signal enrichment and downstream peak calling, resulting in more contiguous peak profiles across centromeres. Karyotype plots were made using KaryoploteR package (version 1.34.2) available in R (Gel & Serra, 2017).

Interestingly, in *A. thaliana*, enrichment peaks were detected prior to kmerRRR based adjustment in regions outside of *CEN180* in chromosome 2 (Fig 4A). Following incorporation of local mappability information, these peaks were no longer observed, despite an increase in total read counts and overall coverage in both treatment and control datasets (Supplemental Fig S9). This pattern suggests that the apparent enrichment may have resulted from disproportionate loss of multi-mapping reads in the control file under conventional filtering thresholds. Whether these represent true biological signals or false-positive peaks arising from multi-mapper handling will require independent experimental validation. In contrast, in chromosome 9 of *H*. sapiens, no peaks were detected (Fig 4) prior to kmerRRR processing after applying irreproducibility discovery rate (IDR) filtering (Li et al., 2011). However, after applying kmerRRR based MAPQL adjustment, reproducible peaks became detectable (Fig 4) within the active *alpha satellite* higher-order repeat (HOR) arrays (Altemose, Logsdon, et al., 2022). Intriguingly, the number and breadth of peaks span nearly the entire alpha satellite HOR arrays, exceeding the estimated occupancy of CENP-A protein within individual centromeres (Altemose, Logsdon, et al., 2022; Altemose, Maslan, et al., 2022). This apparent expansion of peak regions may reflect heterogeneity in CENP-A positioning across HOR arrays as observed in a previous study (Altemose, Maslan, et al., 2022). Because these data are derived from bulk sequencing experiments, the observed signal likely represents the aggregate of all potential CENP-A binding sites across the cell population rather than uniform occupancy across a single centromere. Thus, the broad peak distribution may capture cell-to-cell variability in CENP-A deposition and dynamic binding within *alpha satellite* arrays (Altemose, Maslan, et al., 2022).

By leveraging local mappability, we have shown that kmerRRR facilitates more confident handling of multiple mapping reads, thereby restoring biologically relevant enrichment signals in highly repetitive regions that are otherwise masked by standard global MAPQ filtering.

#### kmerRRR usage in small RNAseq

Repetitive DNA sequences can be a source of RNAs involved in genome regulation (reviewed in (Shatskikh et al., 2020; Wei et al., 2021)). Satellite-derived RNAs are implicated in heterochromatin formation (Wei et al., 2021), genome stability (Fonseca-Carvalho et al., 2024; Goenka et al., 2016; Probst et al., 2010; Valgardsdottir et al., 2008), and dosage compensation (Biswas et al., 2024; Lucchesi & Kuroda, 2015). Some repetitive loci are transcribed and processed into small RNAs, which have roles in heterochromatin formation and TE silencing via the piRNA pathway (Moazed, 2009; Wei et al., 2021). Small RNAseq reads are <100 bp (and often <30 bp), making it difficult to confidently assign reads to their locus of origin, particularly within highly repetitive regions. Accurate locus-specific assignment is essential for understanding which repeat arrays are actively transcribed and for dissecting the functional dynamics of repetitive elements within the genome.

To address these challenges, we developed a pipeline that leverages per-base local mappability inferred from k-mer local to global ratios generated by kmerRRR. Using a companion python script (see documentation), users can generate BED files to define locus-unique regions. These BED files enable extraction of reads from BAM files that overlap locus-unique segments, while discarding reads mapping to non-specific repetitive portions of the locus (Fig 5 and Supplemental Fig S10-S13). Using SAMtools (Danecek et al., 2021), users can then merge filtered BAM files and quantify RNA abundance for downstream analyses or characterize transcriptional profiles of specific repetitive loci. We demonstrate this approach with small RNAseq from *D. melanogaster* (Fig 5 and Supplemental Fig S10; Wei *et al* (2021)). By leveraging local mappability information (using k-mer size of 31), kmerRRR retains ~54-fold more reads at the *Rsp* satDNA locus (4,486 reads) than methods based on only globally unique reads (82 reads; Fig 5), allowing us to identify locations within repetitive DNA loci with evidence for transcription. In addition, we applied kmerRRR to all annotated piRNA clusters in the *D. melanogaster* genome and found that it retained substantially more reads (190,979) than standard methods (i.e. Wei et al. (74,404)), retaining ~2.5-fold more reads. Our exploration of smaller and larger k-mer sizes (21, 31, 61, and 151-mers) had minimal impact on our overall assessment of local mappability (Supplemental Fig S10). Nevertheless, within piRNA clusters, kmerRRR identified globally unique regions that were not detected by the BLAST-based approach of Wei *et al* (2021), enabling the retention of additional reads from these loci and improving their representation in downstream analyses.

**Fig5.**
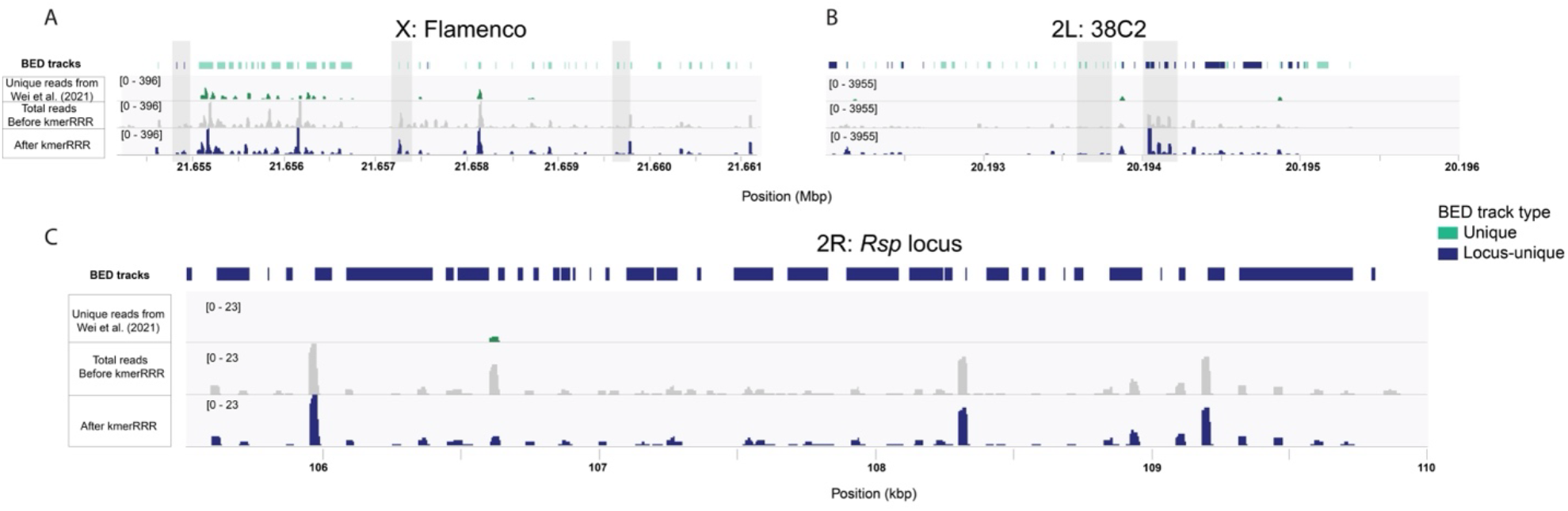
Visualization of LUC-guided read retention of small RNAseq reads. Integrative Genomics Viewer (IGV) illustrate small RNA-seq read coverage across two piRNA clusters, A) flamenco and B) 38C2, and a ~5-kb region of the *Rsp* satDNA locus (Eickbush et al., 2026; Khost et al., 2017) before and after kmerRRR-based filtering. The small RNAseq data are from Wei *et al* (Wei et al., 2021) BED tracks (light green for unique and dark blue for local unique) were generated from kmerRRR using LUC-defined locus-unique and globally unique (unique) regions. Reads overlapping these locus-unique regions were retained in the modified BAM files (dark blue bars), whereas reads not overlapping the BED intervals were excluded (grey bars). Compared to retention strategies based solely on globally unique regions as described by Wei *et al* (2021), kmerRRR retains a greater number of reads by leveraging local mappability information. The plots show alignments from the original alignment output (as described in Wei *et al* (2021); green bars), with grey rectangles indicating reads that are absent in the retained BAM files due to lack of overlap with LUC-defined regions. Overall, this visualization demonstrates that LUC-guided filtering increases informative read retention while preserving locus-specific enrichment patterns.

## Discussion

Here we introduce a new tool, kmerRRR, to improve mappability in repetitive genome regions. kmerRRR infers local mappability by leveraging local-to-global k-mer ratios, enabling the identification of locus-unique characters (LUCs). By distinguishing repetitive regions that are locus-unique from those that are globally repetitive, kmerRRR provides a refined view of genomic mappability, particularly within complex repetitive landscapes such as centromeres, piRNA clusters, and satellite arrays. We found that some repetitive loci (*e*.*g. Rsp* satDNA and 38C2 piRNA cluster of *D. melanogaster, A. thaliana* centromeres) are surprisingly mappable using local mappability, despite consisting of highly repetitive elements.

The concept of local mappability can help retain informative reads that map multiply within, but rarely found outside, a locus. When we apply this local mappability framework to downstream genomic analyses, we circumvent some of the challenges with information loss in repetitive regions with standard quality control workflows. By retaining locus-unique reads in protein-DNA interaction assays such as ChIP-seq and CUT&Tag (Park et al., 2025; Park, 2009), we enhanced signal detection and peak calling in repetitive regions without increasing background noise or false-positive enrichment. We found that peak calling following kmerRRR processing identified signals across broader portions of centromeric higher-order repeats (HOR) arrays than conventional analyses, suggesting heterogenous CENP-A occupancy across HOR units within and among cell populations, consistent with previous observations (Altemose, Maslan, et al., 2022). We also demonstrated the utility of kmerRRR in small RNA sequencing datasets, where locus-unique regions enable confident read assignment in otherwise repetitive contexts. A similar concept has been useful in identifying piRNA clusters in *D. melanogaster* (Chen et al., 2021). Our application of kmerRRR to *D. melanogaster* small RNAseq data revealed transcription of the *Rsp* satDNA locus and piRNA clusters, supporting the production of small RNAs implicated in heterochromatin formation and TE silencing, respectively (Tóth et al., 2016; Wei et al., 2021). Together, these results highlight the broad applicability of the concept of local mappability and kmerRRR workflows across different genomic assays. Our findings suggest that local mappability is a useful framework for extracting functional and regulatory information from repetitive DNA.

K-mer based strategies are widely used for reference free alignment, structural variation discovery, and genome comparison (Moeckel et al., 2024; Voichek & Weigel, 2020). These approaches have been especially useful in analyzing repetitive genome regions across species. For example, k-mer based approaches have been used to create satDNA libraries across species to characterize polymorphisms within satDNA arrays (*e*.*g*. (Wei et al., 2014) (Altemose et al., 2014)). The versatility and computational efficiency of k-mer methodologies make them powerful tools for biological data analysis. kmerRRR extends this paradigm by integrating k-mer derived local mappability directly into genome assemblies. Defining these locus-unique regions can uncover potentially interesting biology. For example, in *A. thaliana*, centromeres on chromosomes 2 and 3 exhibit fewer locus-unique regions compared to other chromosomes, a pattern that may reflect increased interchromosomal exchange or reduced-chromosome specific homogenization of repeats (Fig 2A, 3A). And using locus-unique k-mers to retain informative repetitive reads in functional genomic workflows can improve our inferences about sequence features associated with functionally important domains (e.g. Fig 4 & 5 and Supplemental Fig S9).

However, the utility of kmerRRR depends on several analytical choices, particularly the selection of k-mer size. Different values of *k* can influence the inference of local mappability and the identification of locus-unique regions (Supplemental Fig S1-S3). In general, larger k-mers provide greater sequence specificity and may increase the number of locus-unique positions identified within repetitive loci, thereby increasing the number of reads retained during downstream analyses (Fig 2 and Supplemental Fig S1-S5 & S10-S13). However, the optimal *k* depends on the biological question and sequencing assay. For analyses aimed at characterizing local mappability within repetitive DNA, it may be advantageous to select *k* values that approximate the size of the repeat monomer or to evaluate multiple *k* values to determine which best reflects the underlying repeat architecture. For short-read assays such as small RNAseq, users should carefully evaluate the impact of k-mer size on read retention, as larger k values may substantially increase the number of reads assigned to repetitive loci. Consequently, we recommend exploring multiple *k* values and assessing the biological plausibility of the resulting mappability profiles.

In addition, to parameter selection, kmerRRR relies on a high-quality genome assembly and accurate annotation of the loci of interest. Errors in genome annotation or locus definition may affect local mappability inference and downstream analyses. Furthermore, analyses of large genome, such as the human genome, can require substantial computational resources, including increased memory usage and storage capacity. Many of these challenges are becoming less restrictive as genome assemblies continue to improve in quality and completeness.

Telomere-to-telomere (T2T) genome assemblies are becoming increasingly available, revealing repetitive regions at unprecedented resolution (Li & Durbin, 2024). In this emerging genomic landscape, tools such as kmerRRR will be essential for interpreting and functionally interrogating repetitive arrays. Beyond improving peak calling and transcriptional locus assignment, kmerRRR offers a framework for studying the structural and functional biology of repetitive regions. For instance, LUC informed sequence selection may facilitate the design of locus-specific guide RNAs (Supplemental Fig S7) for targeted manipulation of satDNA arrays using CRISPR-based approaches, enabling experimental dissection of satDNA structure and function across strains and species (Eickbush et al., 2026).

Overall, kmerRRR provides an extensible strategy for leveraging local sequence context to recover biologically meaningful information from repetitive genomic regions that are traditionally masked by conventional mappability constraints.

## Acknowledgments

We thank Logan Edvalson, Leila Lin, and Sreepathi Pai for testing the code and providing feedback. We also thank Justin Fay for providing feedback on the early implementation of kmerRRR. We especially thank John Sproul, Qandeel Zeb, and Emiliano Martí for discussions on earlier implementations of the k-mer ratio. This work was funded by a grant from the National Institutes of Health General Medical Sciences (R35GM119515) to AML. We thank Nathaniel and Helen Wisch for their support.

## Data availability

All data are previously published and publicly available through NCBI’s SRA accessions listed in Table 1. The code and data files required to reproduce the analyses are available in Figshare (DOI forthcoming). kmerRRR is freely available under MIT license in Github: (https://github.com/LarracuenteLab/kmerRRR).

## Supporting information

### Supplementary figures

**Fig. S1.**
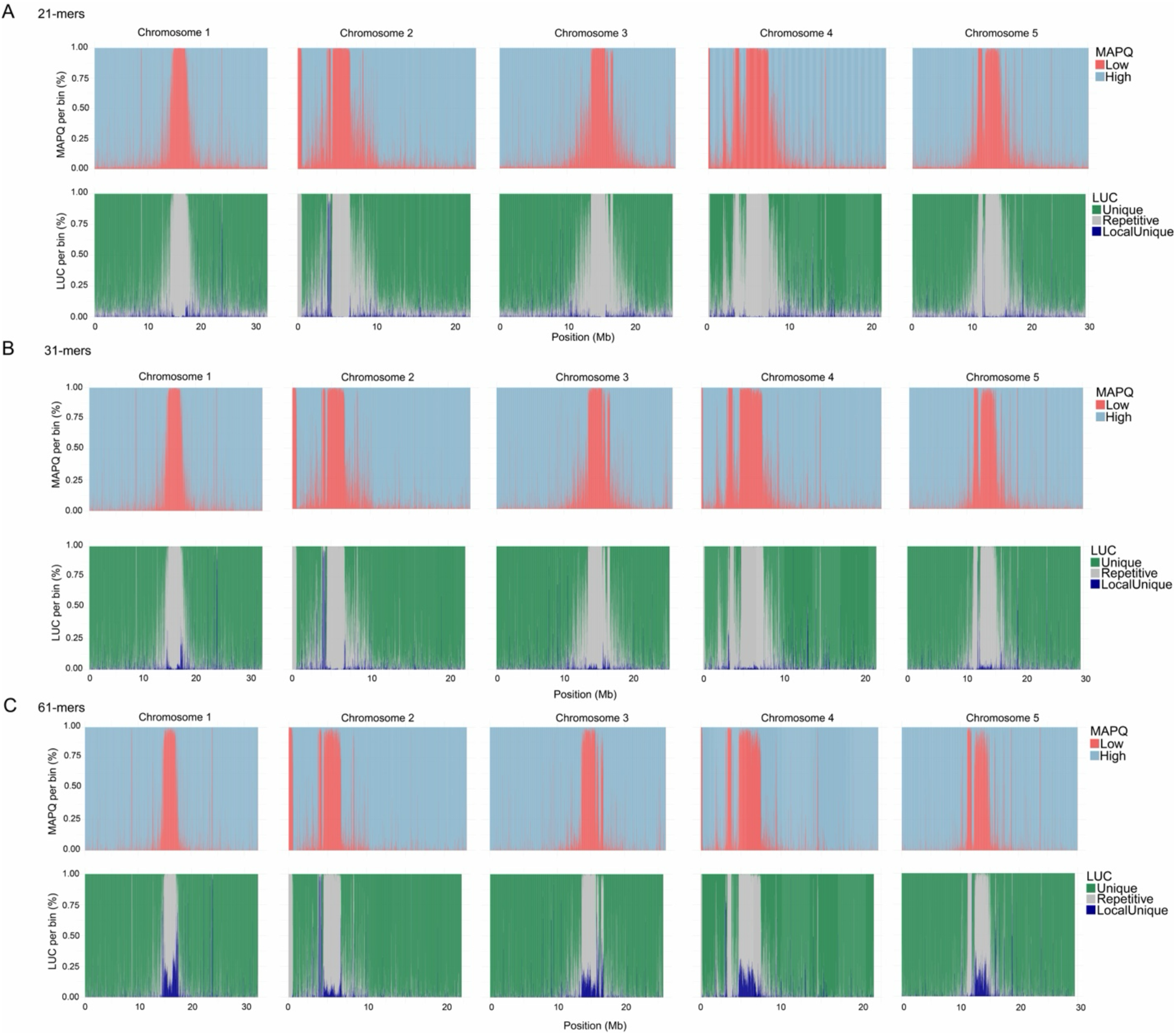
Genome-wide mappability distribution across the near T2T genome of *A. thaliana* for different k-mer lengths. Mappability profiles for k-mer lengths of 21 (A), 31 (B), and 61 (C). The top panels show mappability estimates obtained by remapping synthetic reads of corresponding lengths of *k* across the genome. The bottom panels display Locus-Unique Characters (LUCs) inferred by kmerRRR based on local mappability and local-to-global k-mer frequency ratios at nucleotide resolution for the corresponding k-mer values.

**Fig. S2.**
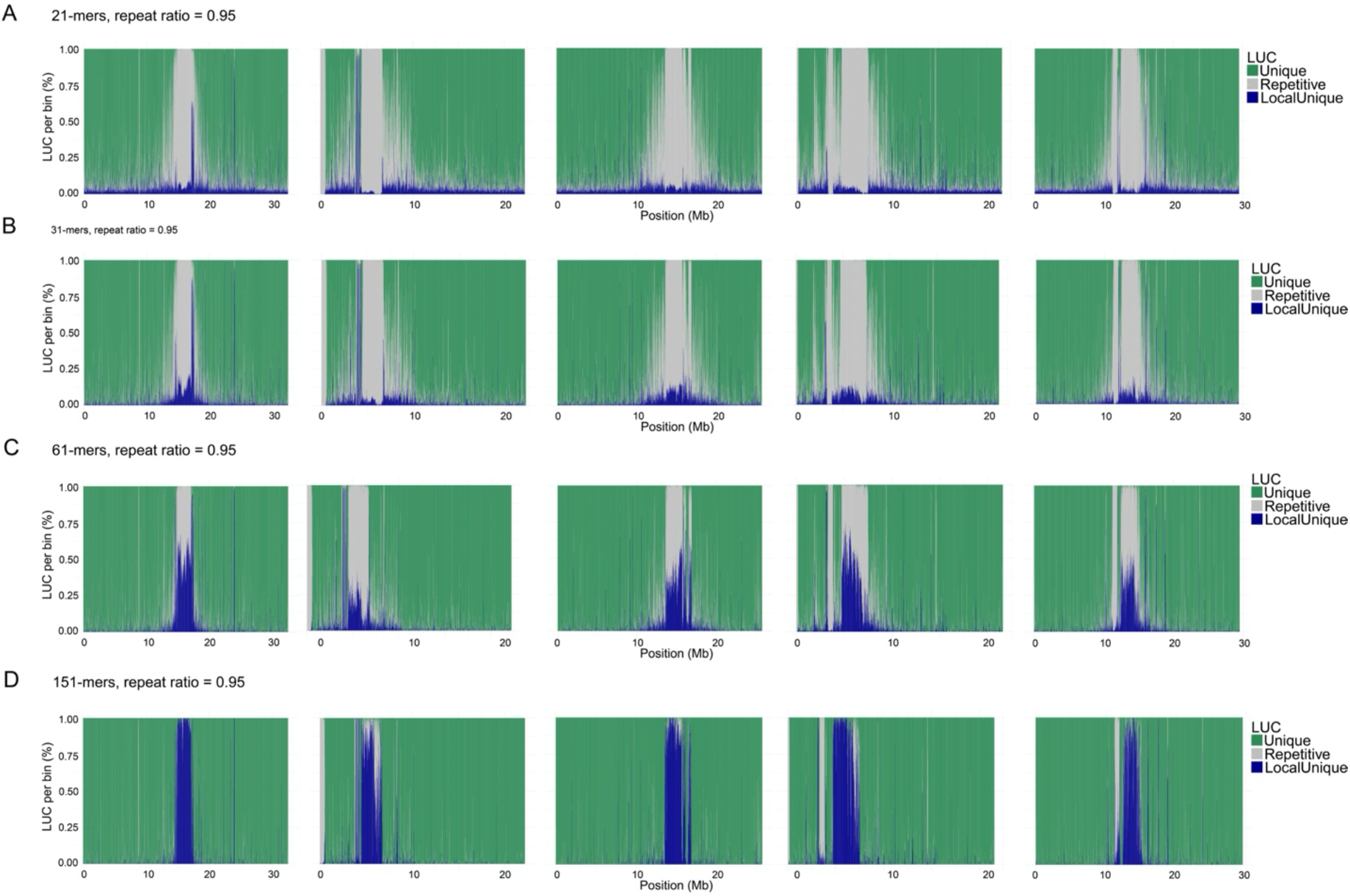
Distribution of Locus-Unique Characters (LUCs) at different k-mer lengths for *A. thaliana* genome. Genome-wide distribution of LUCs inferred by kmerRRR using k-mer lengths of 21 (A), 31 (B), 61 (C), and 151 (D), with a mean repeat ratio threshold of 0.95. Lowering the mean repeat ratio threshold increases the number of regions inferred as locus-unique across loci, however, this increase occurs at the expense of reduced specificity and accuracy across all tested k-mer lengths.

**Fig. S3.**
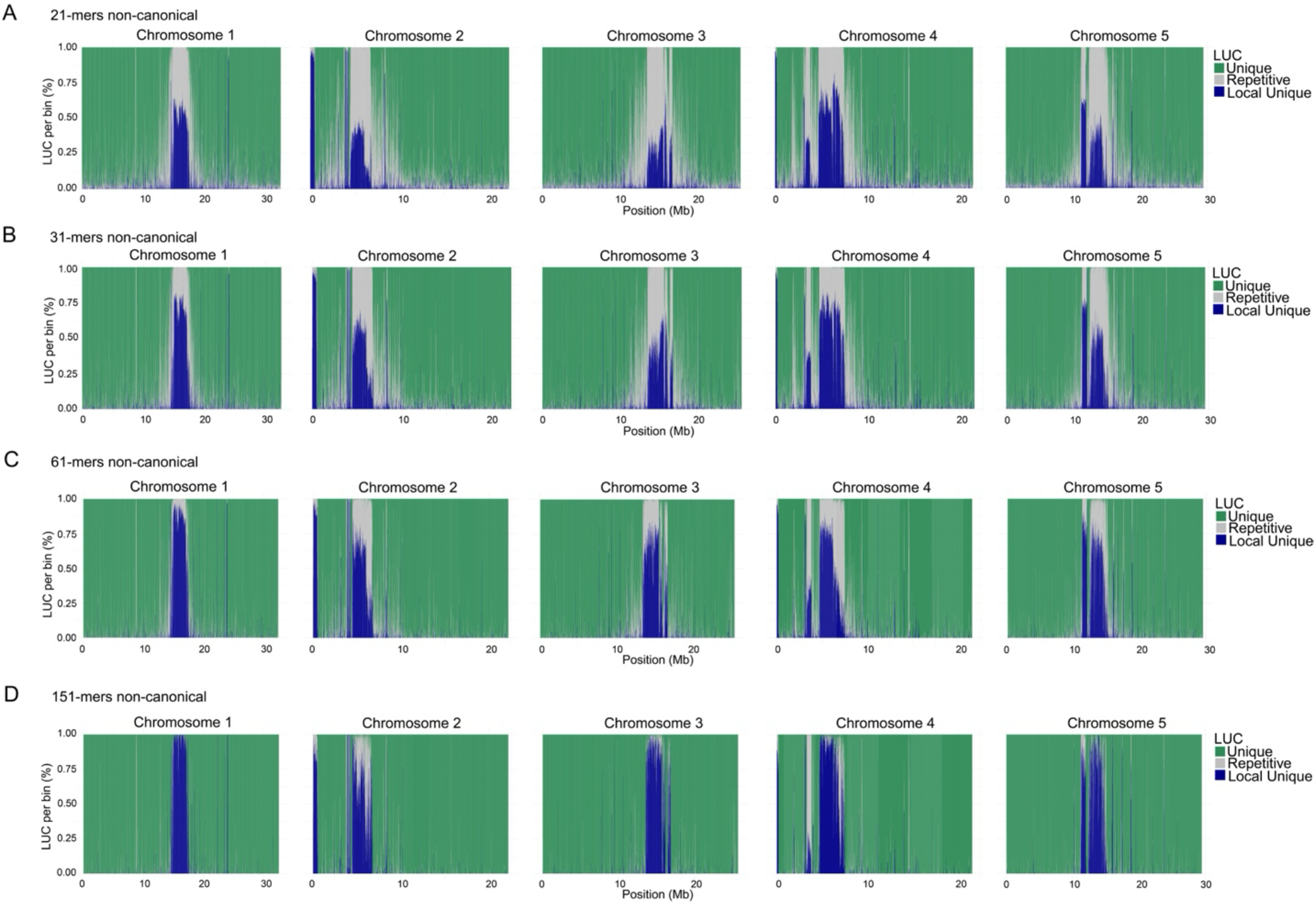
Distribution of Locus-Unique Characters (LUCs) inferred using non-canonical k-mer counting for *A. thaliana*. Genome-wide distribution of LUCs inferred by kmerRRR using non-canonical k-mer counts for k-mer lengths of 21 (A), 31 (B), 61 (C), and 151 (D), with a mean repeat ratio threshold of 1. In the non-canonical framework, k-mers are counted exactly as they occur in the genome assembly without collapsing reverse-complement pairs, thereby preserving strand specificity. Compared with canonical k-mer counting, the non-canonical approach results in substantial differences in LUC inference, with more regions classified as locus-unique despite being globally repetitive under non-canonical counting, particularly for lower values of *k*. This behavior arises because strand-specific sequences are treated as independent k-mers. The use of non-canonical k-mers may therefore be advantageous for strand-sensitive applications such as small RNA sequencing analyses involving the ping–pong amplification cycle or strand-specific RNA-seq pipelines.

**Fig. S4.**
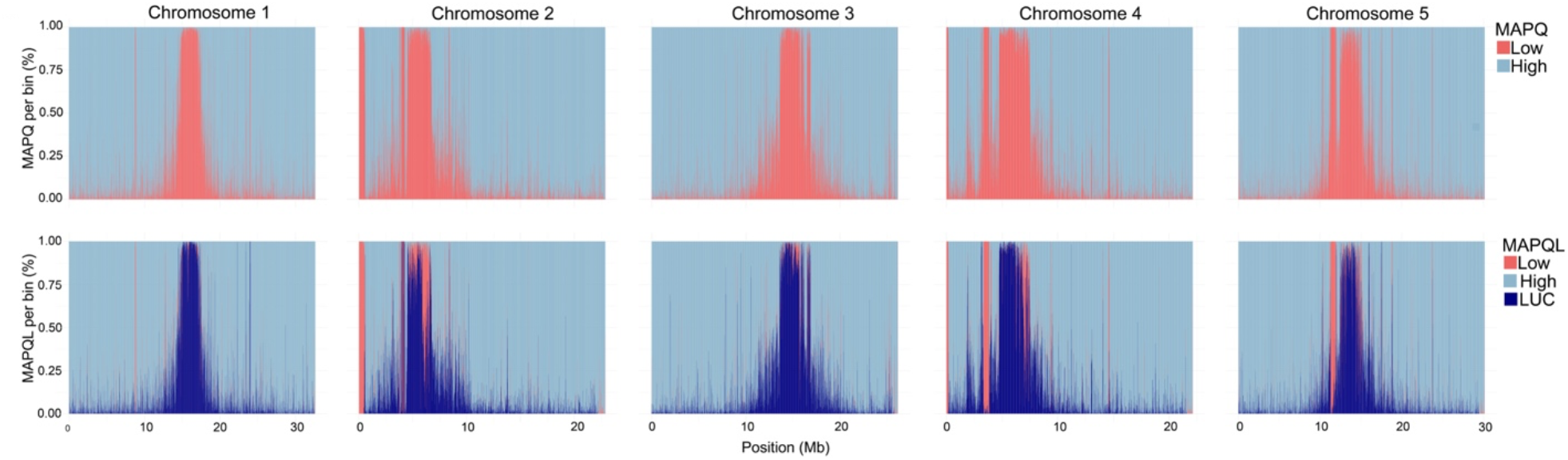
Distribution of LUC-adjusted MAPQL across chromosomes in *A. thaliana*. Genome-wide distribution of reads with LUC-adjusted mapping quality (MAPQL) using a cutoff value of 0.95 across each chromosome for 151-mers. Lowering the cutoff value increases the number of regions where MAPQ scores are adjusted using MAPQL, resulting in a greater number of reads being retained in the BAM files. However, this increase comes with a trade-off between sensitivity and accuracy. At lower cutoff values, more reads are modified even though they originate from regions that are not strictly locus-unique but rather meet the threshold of ≥95% locus-uniqueness.

**Fig. S5.**
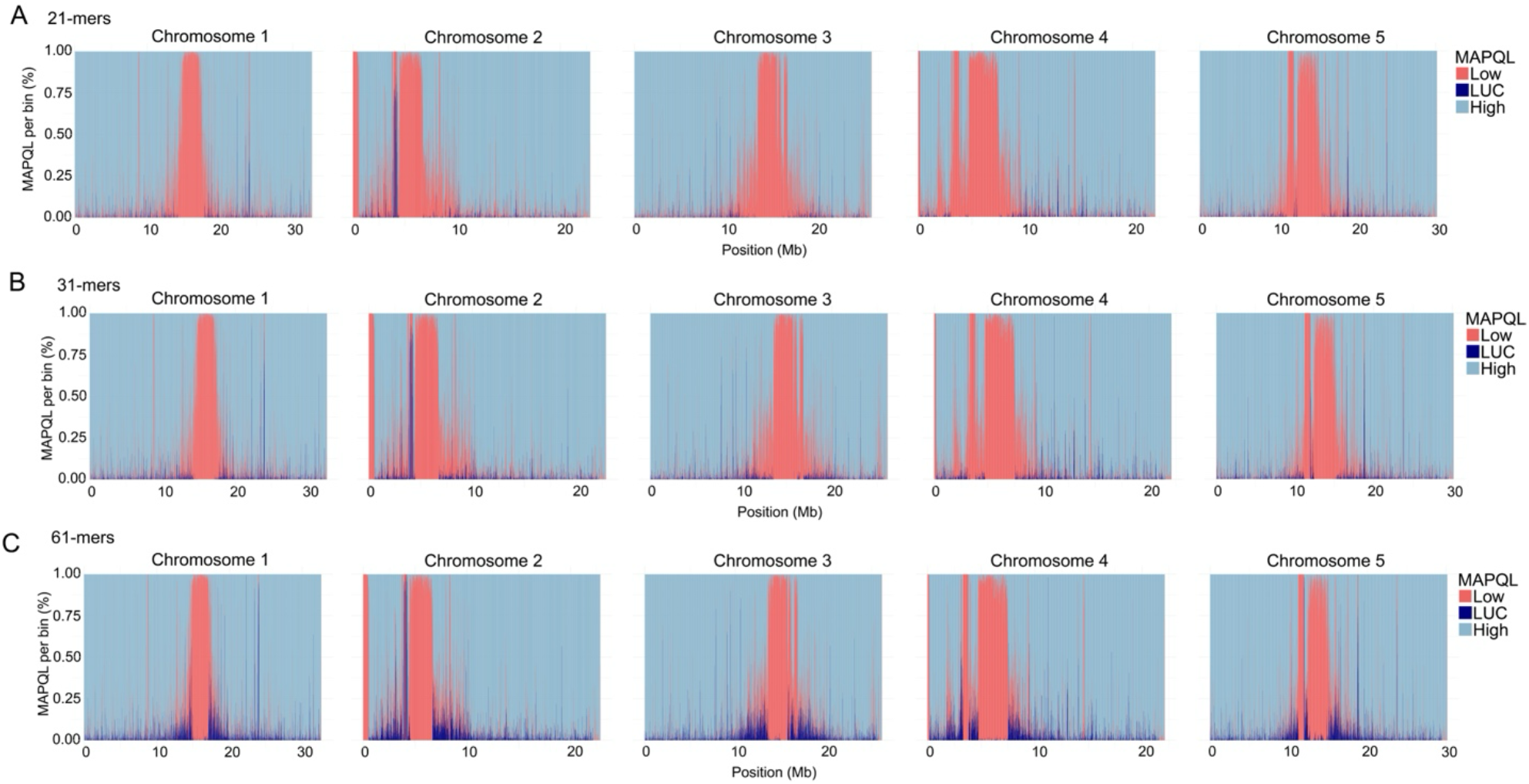
Distribution of MAPQL across chromosomes for different k-mer lengths. Distribution of LUC-adjusted mapping quality (MAPQL) across each chromosome in *A. thaliana* using k-mer lengths of 21 (A), 31 (B), and 61 (C), with a cutoff value of 1. Simulated 150 bp paired-end HiSeq reads generated using ART were used for this analysis. At lower k-mer lengths, kmerRRR has reduced ability to accurately infer locus-unique regions within complex satellite repeats, leading to fewer regions being confidently assigned as locus-unique. Increasing the k-mer length improves the resolution of local mappability and enables better identification of locus-unique regions within repetitive genomic contexts.

**Fig. S6.**
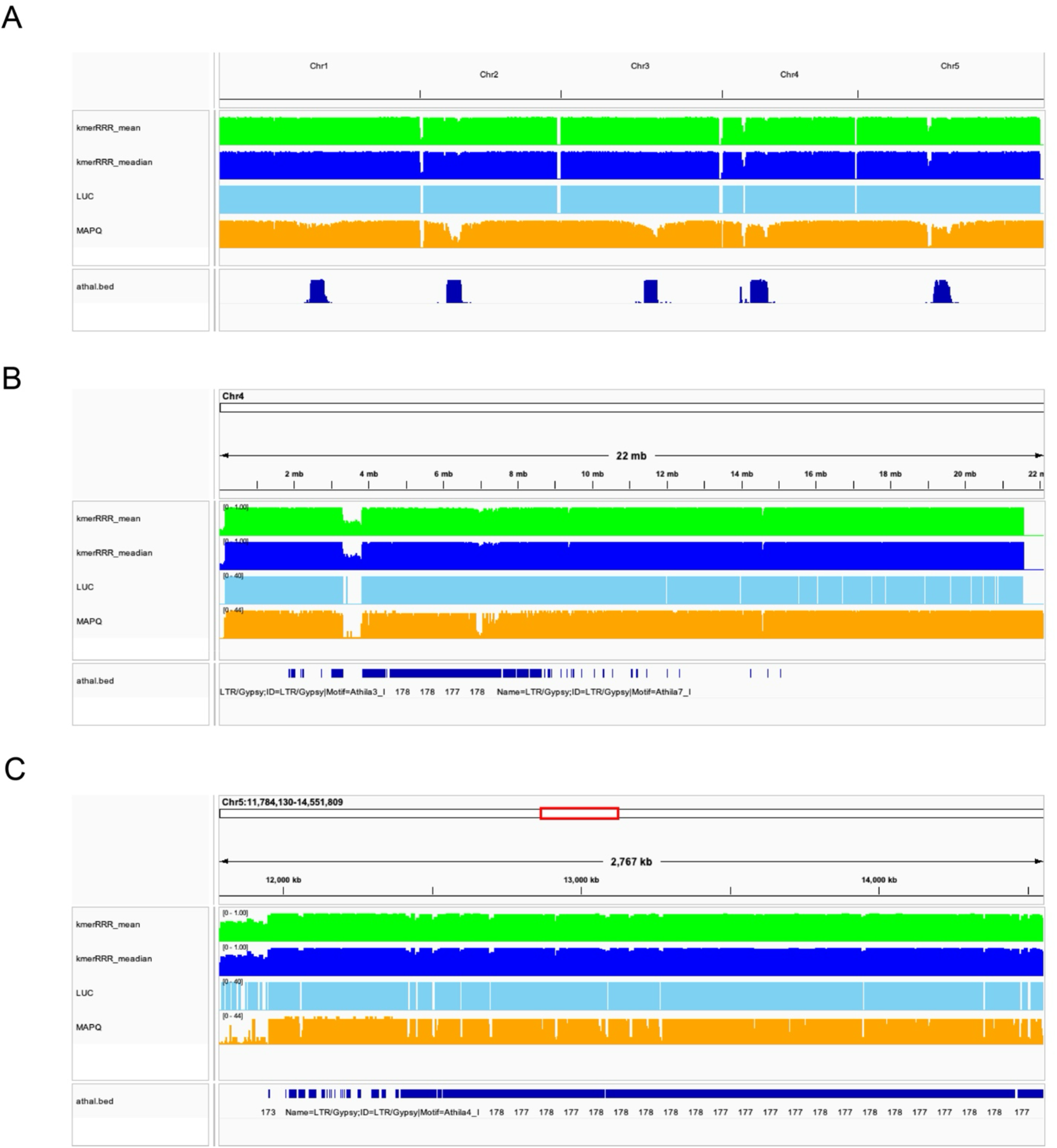
IGV snapshots showing genome-wide and locus-specific distribution of LUCs in *A. thaliana*. IGV snapshots of the *A. thaliana* genome showing (A) the whole genome view, (B) chromosome 4, and (C) a zoomed in view of the centromere of chromosome 5. The bottom annotation track indicates the centromeric island regions. kmerRRR can generate bedGraph outputs based on per-base LUC inference, including the mean (green bars) and median k-mer (blue bars) ratio at the nucleotide level. These tracks allow visualization of the distribution of locus-unique characters (LUCs) (light blue bars) across specific genomic loci, along with mapping quality (MAPQ) (orange bars) values when available.

**Fig. S7.**
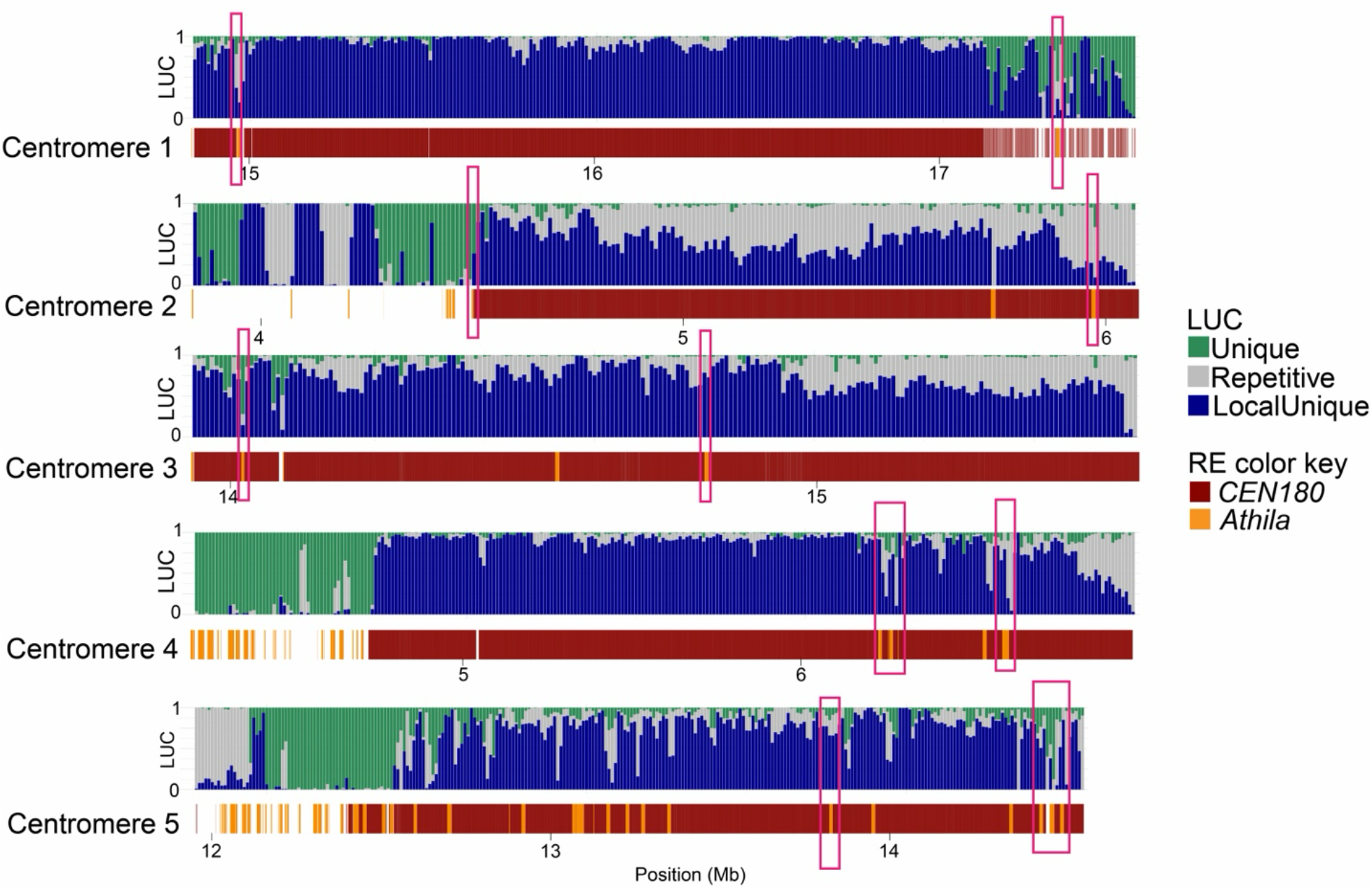
Distribution of locus-unique characters (LUCs) across the centromeric islands of each chromosome in *A. thaliana*. Pink rectangles highlight regions of globally unique sequence at the junctions between transposable elements (TEs) and *CEN180* satellite repeats. These junctions represent globally unique sequence contexts within otherwise repetitive regions, illustrating how kmerRRR can be used to identify candidate loci for designing guides or primers for downstream molecular analyses. Karyotype plots were made using KaryoploteR package (version 1.34.2) {Gel, 2017 #80}.

**Fig. S8.**
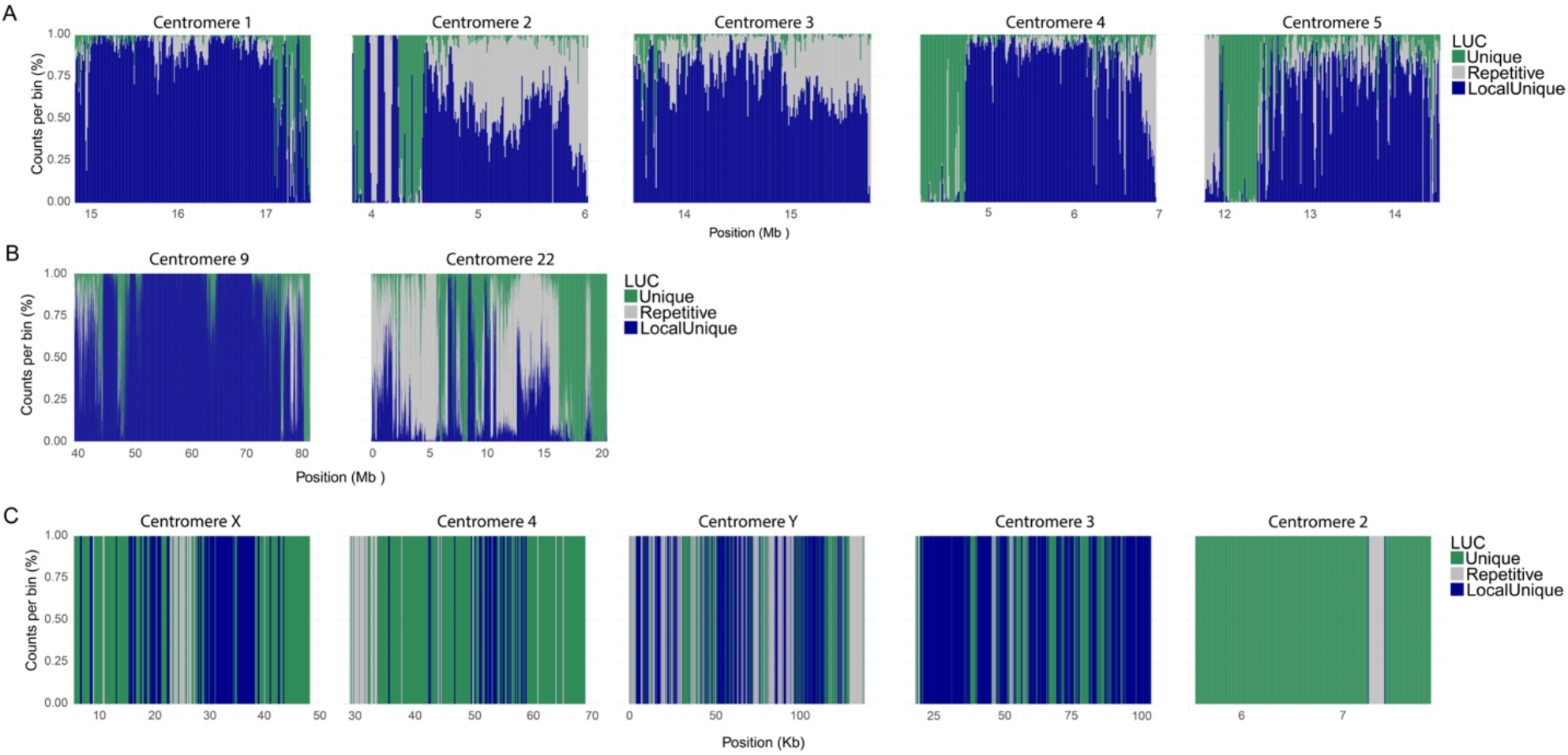
Distribution of LUCs across centromeric islands in different species. Distribution of locus-unique characters (LUCs) across centromeric islands in the genomes of *A. thaliana* (A), *H. sapiens* (B), and *D. melanogaster* (C). This comparison highlights how locus-unique regions are distributed within centromeric repetitive DNA across diverse eukaryotic genomes.

**Fig. S9.**
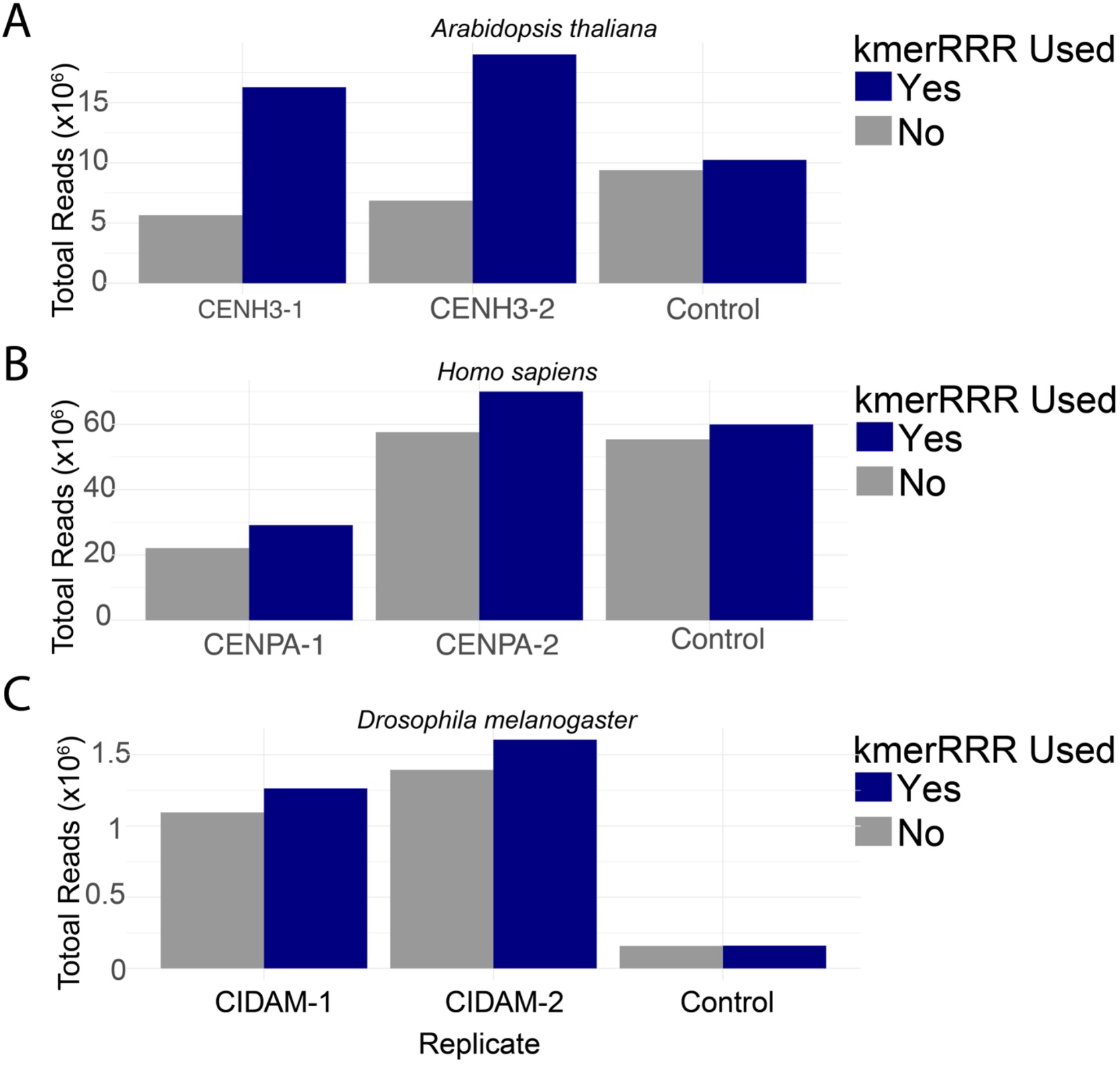
Read retention before and after kmerRRR adjustment with MAPQL. Number of reads in BAM files before and after applying kmerRRR followed by filtering with a mapping quality (MAPQ) threshold of 30. The use of kmerRRR adjusts mapping quality scores using MAPQL based on locally inferred mappability, allowing retention of reads that would otherwise be discarded due to ambiguous mapping. This results in increased read retention in both treatment and control datasets, improving the potential for enrichment analyses during peak calling in centromeric regions of *A. thaliana* (A), *H. sapiens* (B), and *D. melanogaster* (C).

**Fig. S10.**
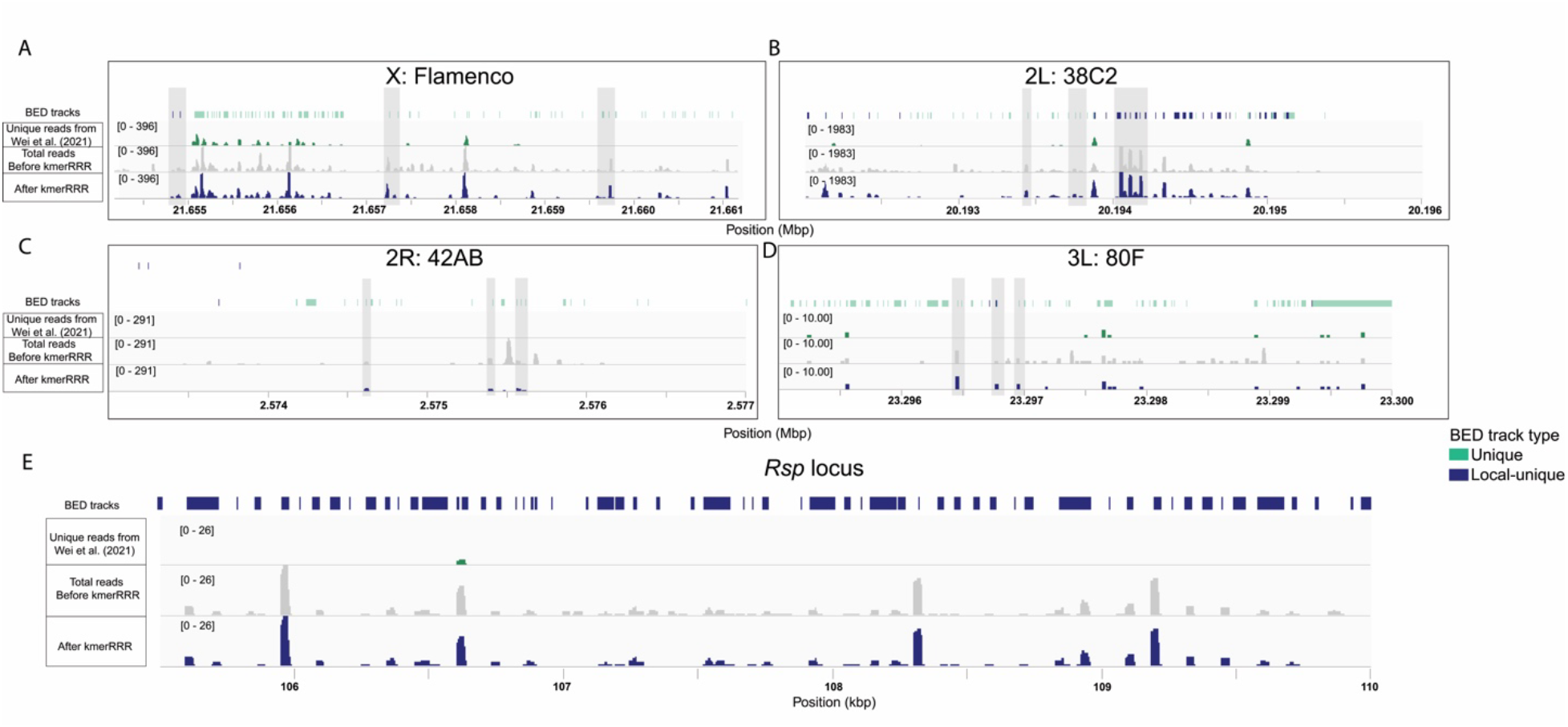
Visualization of LUC-guided read retention of small RNAseq reads with 21-mers. Integrative Genomics Viewer (IGV) illustrate small RNA-seq read coverage across *D. melanogaster* piRNA clusters, A) flamenco, B) 38C2, C) 42AB, ands D) 80F, and a 5kb region of the *Rsp* satDNA locus before and after kmerRRR-based filtering. The small RNAseq data are from Wei *et al* (2021) BED tracks (light green for unique and dark blue for locus-unique) were generated from kmerRRR using LUC-defined locus-unique and unique regions. Reads overlapping these locus-unique regions were retained in the modified BAM files (dark blue bars), whereas reads not overlapping the BED intervals were excluded (grey bars). Compared to retention strategies based solely on globally unique regions as described by Wei *et al* (2021), kmerRRR retains a greater number of reads by leveraging local mappability information (1,90,979 reads for piRNA cluster and 3,764 reads for *Rsp* locus). The plots show alignments from the original alignment output (as described in Wei *et al* (2021); green bars), with grey rectangles indicating reads that are absent in the retained BAM files due to lack of overlap with LUC-defined regions. Overall, this visualization demonstrates that LUC-guided filtering increases informative read retention while preserving locus-specific enrichment patterns.

**Fig. S11.**
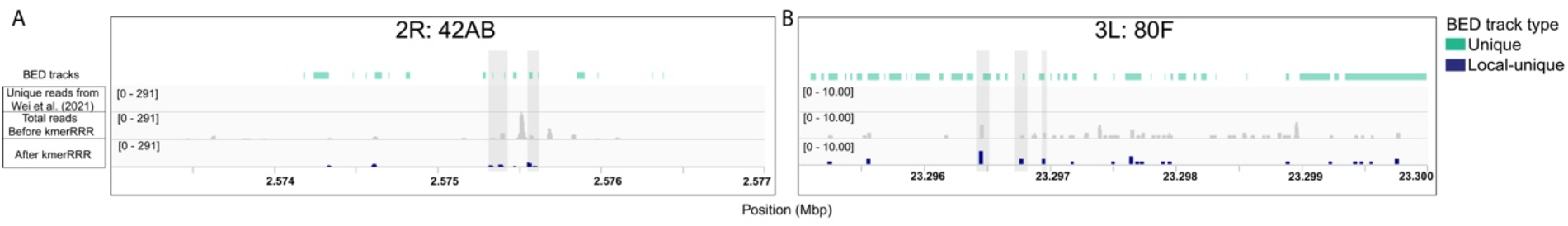
Visualization of LUC-guided read retention of small RNAseq reads with 31-mers. Integrative Genomics Viewer (IGV) illustrate small RNA-seq read coverage across *D. melanogaster* piRNA clusters, A) 42AB, ands B) 80F before and after kmerRRR-based filtering. The small RNAseq data are from Wei *et al* (2021) BED tracks (light green for unique and dark blue for locus-unique) were generated from kmerRRR using LUC-defined locus-unique and unique regions. Reads overlapping these locus-unique regions were retained in the modified BAM files (dark blue bars), whereas reads not overlapping the BED intervals were excluded (grey bars). Compared to retention strategies based solely on globally unique regions as described by Wei *et al* (2021), kmerRRR retains a greater number of reads by leveraging local mappability information. The plots show alignments from the original alignment output (as described in Wei *et al* (2021); green bars), with grey rectangles indicating reads that are absent in the retained BAM files due to lack of overlap with LUC-defined regions. Overall, this visualization demonstrates that LUC-guided filtering increases informative read retention while preserving locus-specific enrichment patterns.

**Fig. S12.**
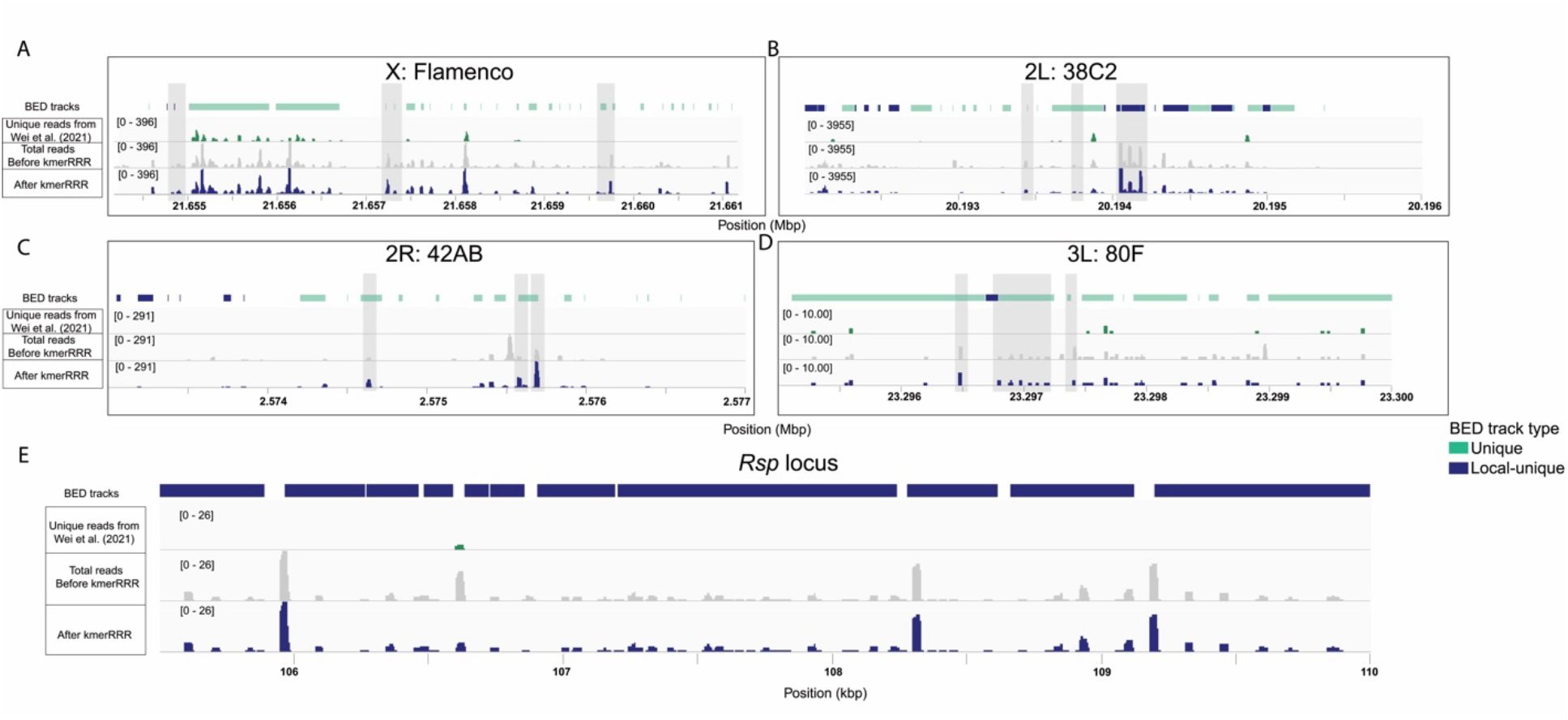
Visualization of LUC-guided read retention of small RNAseq reads with 61-mers. Integrative Genomics Viewer (IGV) illustrate small RNA-seq read coverage across *D. melanogaster* piRNA clusters, A) flamenco, B) 38C2, C) 42AB, ands D) 80F, and a 5kb region of the *Rsp* satDNA locus before and after kmerRRR-based filtering. The small RNAseq data are from Wei *et al* (2021) BED tracks (light green for unique and dark blue for locus-unique) were generated from kmerRRR using LUC-defined locus-unique and unique regions. Reads overlapping these locus-unique regions were retained in the modified BAM files (dark blue bars), whereas reads not overlapping the BED intervals were excluded (grey bars). Compared to retention strategies based solely on globally unique regions as described by Wei *et al* (2021), kmerRRR retains a greater number of reads by leveraging local mappability information (2,44,307 reads for piRNA cluster and 5,847 reads for *Rsp* locus). The plots show alignments from the original alignment output (as described in Wei *et al* (2021); green bars), with grey rectangles indicating reads that are absent in the retained BAM files due to lack of overlap with LUC-defined regions. Overall, this visualization demonstrates that LUC-guided filtering increases informative read retention while preserving locus-specific enrichment patterns.

**Fig. S13.**
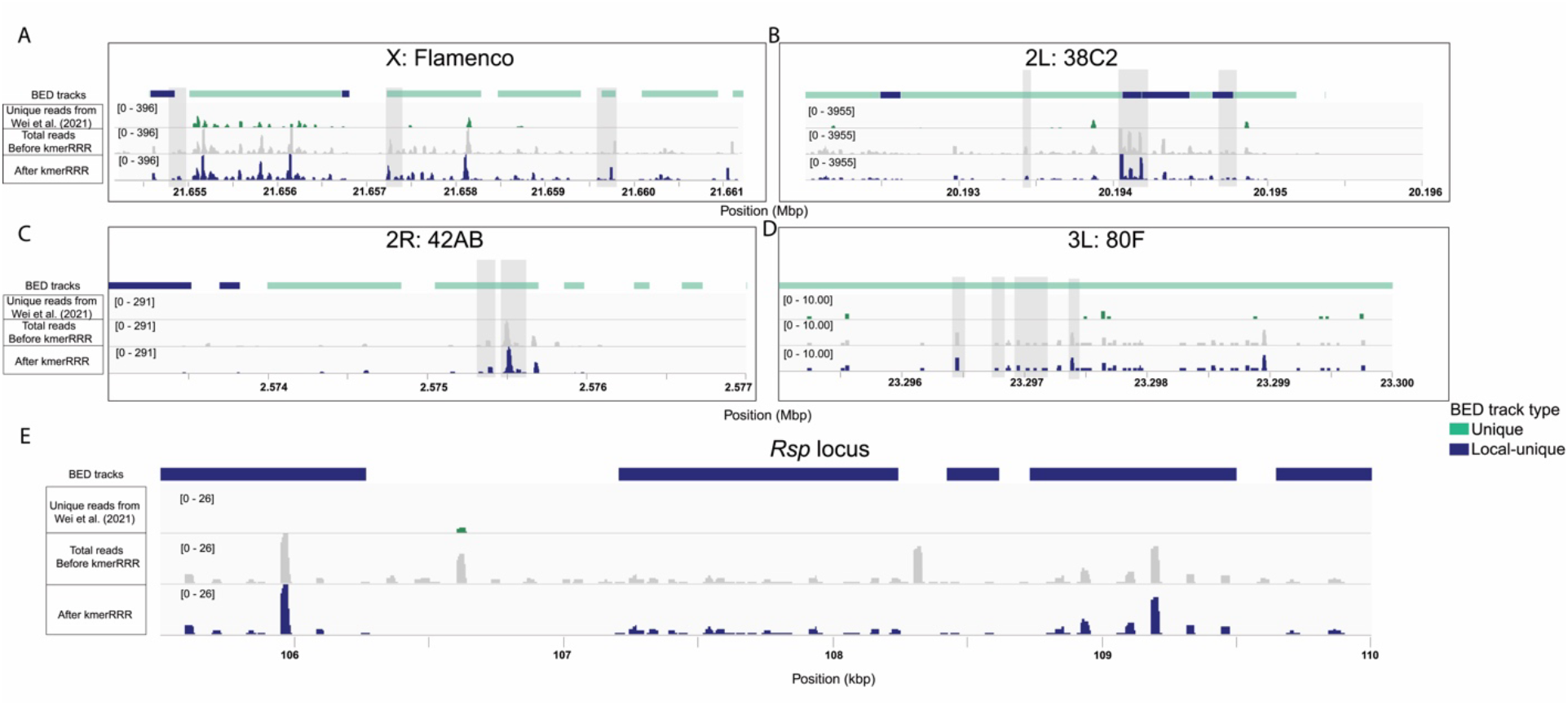
Visualization of LUC-guided read retention of small RNAseq reads with 151-mers. Integrative Genomics Viewer (IGV) illustrate small RNA-seq read coverage across *D. melanogaster* piRNA clusters, A) flamenco, B) 38C2, C) 42AB, ands D) 80F, and a 5kb region of the *Rsp* satDNA locus before and after kmerRRR-based filtering. The small RNAseq data are from Wei *et al* (2021) BED tracks (light green for unique and dark blue for locus-unique) were generated from kmerRRR using LUC-defined locus-unique and unique regions. Reads overlapping these locus-unique regions were retained in the modified BAM files (dark blue bars), whereas reads not overlapping the BED intervals were excluded (grey bars). Compared to retention strategies based solely on globally unique regions as described by Wei *et al* (2021), kmerRRR retains a greater number of reads by leveraging local mappability information (2,74,900 reads for piRNA cluster and 7,492 reads for *Rsp* locus). The plots show alignments from the original alignment output (as described in Wei *et al* (2021); green bars), with grey rectangles indicating reads that are absent in the retained BAM files due to lack of overlap with LUC-defined regions. Overall, this visualization demonstrates that LUC-guided filtering increases informative read retention while preserving locus-specific enrichment patterns.

